# High-Throughput DNA melt measurements enable improved models of DNA folding thermodynamics

**DOI:** 10.1101/2024.01.08.574731

**Authors:** Yuxi Ke, Eesha Sharma, Hannah K. Wayment-Steele, Winston R. Becker, Anthony Ho, Emil Marklund, William J. Greenleaf

**Affiliations:** Department of Bioengineering, Stanford University, Stanford, CA 94305, USA; Department of Genetics, Stanford University School of Medicine, Stanford, CA 94305, USA; Lamar Health, Palo Alto, CA 94305, USA; Department of Chemistry, Stanford University, Stanford, CA 94305, USA; Program in Biophysics, Stanford University, Stanford, CA 94305, USA; Science for Life Laboratory, Department of Biochemistry and Biophysics, Stockholm University, 17165, Solna, Sweden; Department of Applied Physics, Stanford University, Stanford, CA 94305, USA

## Abstract

DNA folding thermodynamics are central to many biological processes and biotechnological applications involving base-pairing. Current methods for predicting stability from DNA sequence use nearest-neighbor models that struggle to accurately capture the diverse sequence-dependency of elements other than Watson-Crick base pairs, likely due to insufficient experimental data. We introduce a massively parallel method, Array Melt, that uses fluorescence-based quenching signals to measure equilibrium stability of millions of DNA hairpins simultaneously on a repurposed Illumina sequencing flow cell. By leveraging this dataset of 27,732 sequences with two-state melting behavior, we derived a refined NUPACK-compatible nearest-neighbor model, a richer parameterization nearest-neighbor model that exhibits higher accuracy, and a graph neural network (GNN) model that identifies relevant interactions within DNA beyond nearest neighbors. All models provide improved accuracy in predicting DNA folding thermodynamics, providing improvements relevant for *in silico* design of qPCR primers, oligo hybridization probes, and DNA origami.

## Introduction

The thermodynamics of DNA secondary structure formation is fundamental to understanding diverse biological processes, such as DNA replication^1^ and repair^2^, and is widely used in biotechnological applications, including the design of PCR primers^3^, the engineering of in-situ hybridization (ISH) probes^4^, the design of guide sequences for genome editing tools^5^, and for DNA nanotechnology^6,7^. Numerous algorithms have been developed to predict DNA secondary structure thermodynamics, many of which rely on nearest-neighbor models^8–10^.

Briefly, nearest-neighbor models assume that the total folding energy of DNA can be calculated by summing up the energies of each pair of neighboring base pairs. With this assumption, the “folded state” can be defined as the minimum free energy (MFE) base pairing configuration, identified using dynamic programming^11^. Additionally, by using partition functions, it is possible to calculate probabilities of the DNA in alternative secondary structures based on these computed free energies. However, nearest-neighbor models often struggle to accurately capture the sheer diversity and complexity of DNA secondary structure motifs, including hairpin loops, mismatches, and bulges. One reason for this limitation is the lack of experimental data spanning those loop features upon which these models are built, leading to inaccuracies in prediction. For example, the most widely used parameter set from *SantaLucia et al. 2004* used data from 108 sequences to derive 12 parameters for Watson-Crick (WC) base pairs, and 174 sequences to derive 44 parameters for internal single mismatches^8^.

This data bottleneck has been due to the laborious nature of optical melting^12^ and differential scanning calorimetry^13^ experiments, traditionally considered the gold standards of DNA secondary structure dynamics. Several groups have attempted to overcome this bottleneck, usually with fluorescent melting in bulk solutions in well plates^14–17^, reaching a throughput on the scale of thousands. Still, due to the extremely large combinatorial DNA sequence space, these works are limited to a specific type of DNA structural element in a fixed sequence scaffold.

To address this data generation bottleneck, we developed a method for systematic, accurate, high-throughput measurements of nucleic acid secondary-structure motif thermal stability, enabling large-scale, quantitative measurements of the thermodynamics of nucleic acid secondary structure. We demonstrate that nearest-neighbor thermodynamic parameters inferred from these data are nearly identical to those observed from bulk thermal melting experiments for WC base pairs but are improved for loop sequence motifs. The improved nearest-neighbor model parameter sets are capable of accurately predicting DNA duplex folding energy on orthogonal unseen data sets. Furthermore, by deploying advanced computational methods including deep learning to model DNA thermodynamics, we develop a state-of-the-art thermodynamic model for DNA hairpin folding with accuracy comparable to measurement uncertainties.

## Results

### Array melt: A high-throughput technique for quantifying the melting behavior of nucleic acids

We developed the Array Melt technique, a method that allows for the rapid and comprehensive quantification of DNA folding thermodynamics in high throughput. This technique is based on an Illumina MiSeq chip repurposed for high-throughput measurements^18,19^. To assay the melting behavior of a nucleic acid construct, we engineered a common region for annealing a 3’-fluorophore-labeled oligonucleotide to the 5’-end of a hairpin to be investigated. Similarly, we designed another region for annealing a 5’-quencher-labeled oligonucleotide to the 3’-end of the hairpin (**Fig. 1a**). After determining the position and sequence of the variant on the MiSeq chip through sequencing, we annealed Cy3-labeled and Black Hole Quencher (BHQ)-labeled oligonucleotides to these respective binding sites. These annealed oligos have predicted melting temperatures of 74°C, much higher than the highest temperature in our experiments (**Table S1**). At room temperature, this fluorophore-quencher pair results in low levels of fluorescence in each cluster, as the bulk of our hairpin library is folded at room temperature. As the sequence-variable hairpin regions are exposed to increasing temperatures between 20 °C to 60 °C, the distance between Cy3 and BHQ increases, leading to brighter fluorescence signals (**Fig. 1a-b**). Using this measurement scheme, we synthesized a comprehensive library of 42,177 unique hairpin sequences for measurement (**Fig. 1c**). These sequences integrated diverse structural elements into a hairpin scaffold, including Watson-Crick pairing (WC), mismatches, bulges, hairpin loops of various lengths, and other control sequences, including super-stable stem, polynucleotide repeats, and single-strand fluorescence controls (see **Fig. 1c** and Methods).

**Figure 1.**
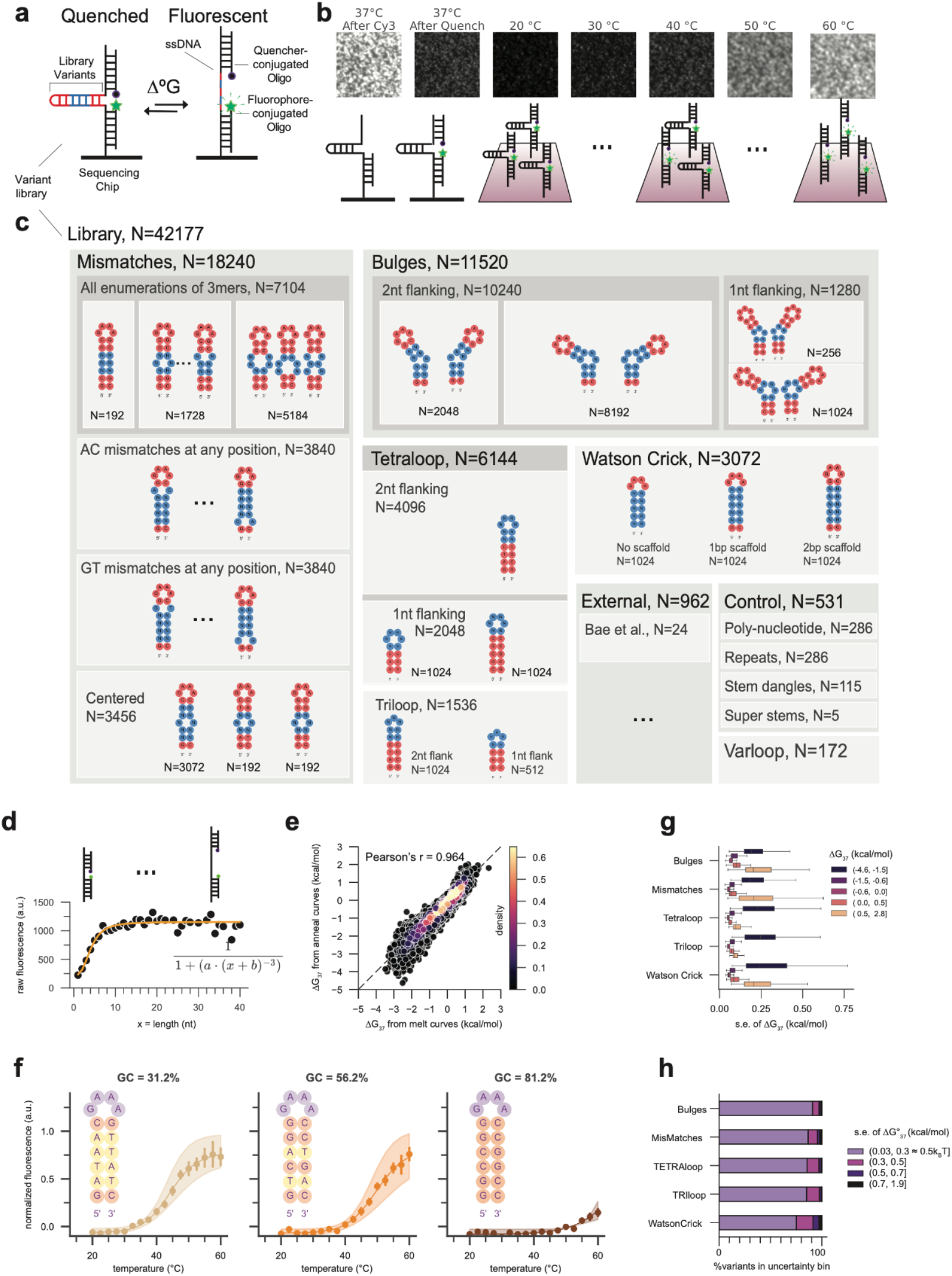
High-Throughput thermodynamic measurement of DNA secondary structures using a fluorophore– quencher system. a. Schematic representation of DNA molecules in a folded (quenched) and an unfolded (fluorescent) state. b. Images of fluorescent DNA clusters immobilized on the sequencing chip surface. Top: Image with only fluorophore-conjugated oligo (Cy3), and with both fluorophore- and quencher-conjugated oligo at increasing temperatures. Bottom: Schematics of DNA molecules in each image. All images are normalized to super-stable stem and repeat control variants for temperature-dependent effects on fluorescence and quenching during the experiment, as shown in **Fig. S1d**. c. Design of library variants between the constant-sequence binding sites for fluorophore- and quencher-conjugated oligonucleotides. Red represents scaffold nucleotides constant within each type, and blue represents the variable nucleotides systematically permuted (‘N’). Numbers under each class indicate the number of unique sequences in each class. d. Fluorescence measurement of control constructs where the single-strand distance between the fluorophore and the quencher increases in single nucleotide steps. The orange line shows the theoretical fit. e. Correlation of ΔG_37_ derived from library variants under increasing temperatures (melt curve, x-axis) and decreasing temperatures (annealing curve, y-axis) f. Representative examples of melt curves for constructs that vary in GC content. g. Pearson correlation coefficient (*R*) of ΔG_37_ for replicates of three melt and one annealing curve experiment. h. Standard error of ΔG_37_ for various construct classes as a function of ΔG_37_.

We performed thorough quality control on the dataset by first requiring variants to accurately fit a two-state model with melting behavior observable within our temperature measurement range (see Methods for our filtering strategy). Total variants that pass these stringent thresholds include 6,393,050 individual melt curves (including technical replicates) spanning 27,732 sequence variants for subsequent analysis with a standard two-state melt model.

### Sensitivity and precision of the Array Melt method

To calibrate the amount of fluorescence observed as a function of the distance between Cy3 and BHQ, we evaluated the fluorescence signals of a series of repeat sequence variants. These variants were designed to systematically increase the distance between the fluorophore and quencher by incorporating increasing numbers of mono-, di- or tri-nucleotide repeats. The fluorescent signals, aggregated over repeats of the same length, showed a nearly linear response to distance variations up to approximately 8 nucleotides (**Fig. 1d**). Our observed length-fluorescence curve closely aligns with a theoretical static quenching curve (Methods). We normalized the fluorescent signals using unfolded and folded controls designed within the library (**Fig. S1b**). Three example melt curves in **Fig. 1f** illustrated increasing observed melting temperatures and increasing GC content of the variable region. Technical replicate measurements from Array Melt data showed high correlation, with *R* > 0.94 (**Fig. S1c**). Melt curves were also highly correlated with anneal curves (*R* = 0.964), confirming that the DNA molecules were measured at equilibrium (**Fig. 1e**). Upon analyzing our measurement precision (**Methods**), we found that nearly 90% of the tested variants displayed Δ*G*_37_ uncertainties within 0.3 kcal/mol, or 0.5 kBT (**Fig. 1h)**. Overall, these data demonstrate that the equilibrium, aggregate fluorescence signals obtained from the Array Melt method provide highly accurate melting information and can serve as a “molecular ruler” capable of resolving single nucleotide distance variations.

### UV melting validation

To further validate our method, we compared the results from the Array Melt method to those obtained using conventional UV melting methods on a cloud lab platform. We selected random representative variants from each category within the Array Melt library and sent the synthesized oligonucleotides to Emerald Cloud Lab (ECL) for analysis. Using the variant information and UV melting experiment parameters as input, scripts written in Symbolic Lab Language (SLL) automatically generated experiments as ECL protocol objects. These protocols were then queued for remote execution in a physical laboratory. Raw melting curve data from ECL were automatically analyzed by custom Python code, and samples that failed to pass quality control were resubmitted to ECL for re-measurement (**Fig. 2a, Fig. S2a, Methods**). A comparison of the melting profiles of random library variants showed good agreement between the two methods (**Fig. 2c-e**, R=0.85 for ΔG_37_, or R^2^= 0.62 after the linear adjustment described later). The melting temperature was found to be independent of DNA oligo concentration (**Fig. S2b**), suggesting intramolecular folding. Notably, we observed a systematic offset between the UV measurements and our Array Melt results, which could be due to the presence of the hairpins on a flow-cell surface, or the additional sequence elements included to facilitate our quenching-based readout. To account for these potential systematic differences, we applied a single offset to the melting points observed in our dataset and used the corrected results for subsequent analysis (**Methods**; on average, a correction of 9.35°C in Tm or approximately 0.7 kcal/mol in ΔG_37_ was applied). Compared to standard nearest-neighbor model predictions from NUPACK software, the measured values showed moderate improvement for ΔG and Tm, and considerable improvement for ΔH (**Fig. 2e, Fig. S2c**). This UV melting validation demonstrated the ability of the Array Melt method to accurately measure the folding energies of DNA and generalize to in-solution systems.

**Figure 2.**
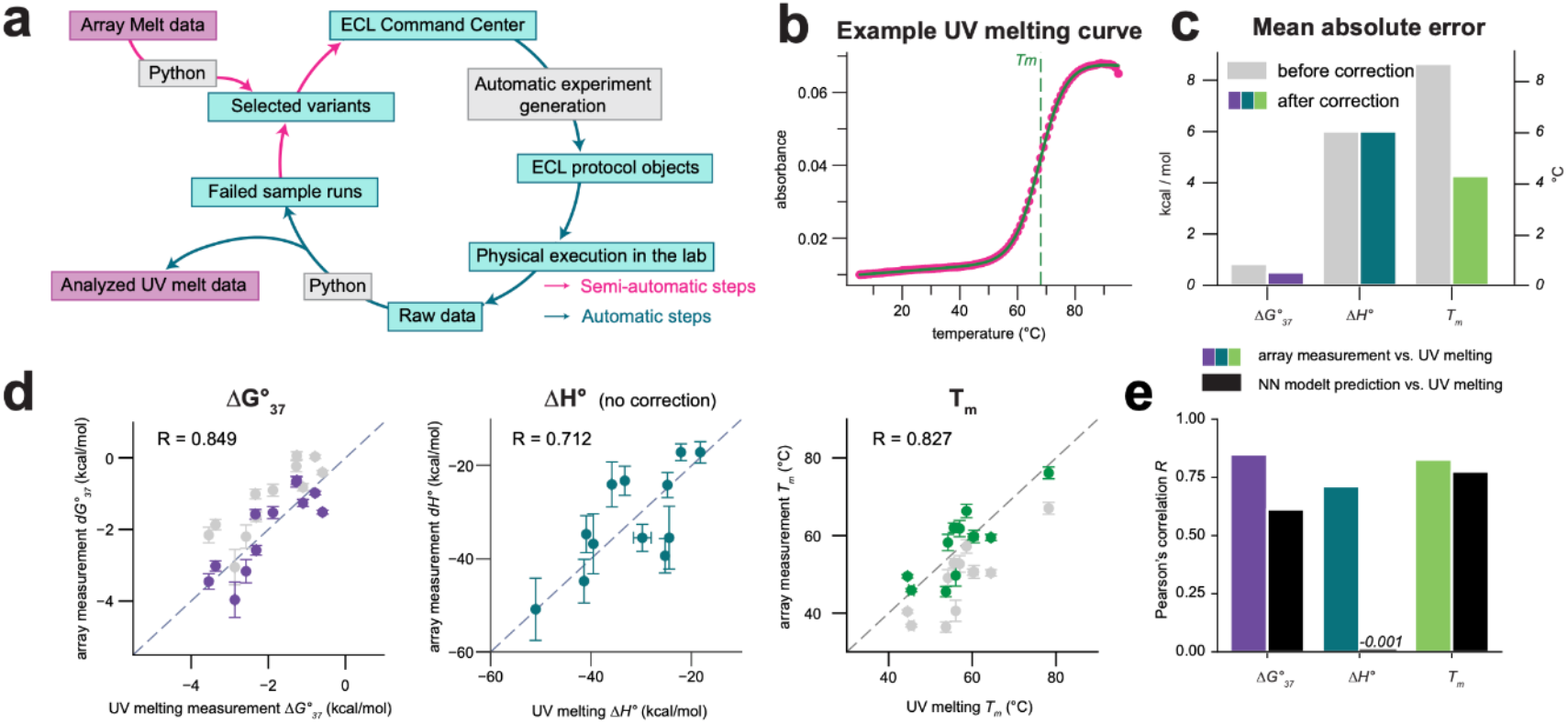
Cloud lab UV melting validation of the method. a. Schematic representation of UV melting data collection and analysis using the Emerald Cloud Lab (ECL) platform. Magenta arrows indicate semi-automatic steps, and teal arrows indicate automatic steps. b. Example UV melting curve of the variant WC2318. Pink dots represent measured data points of the cooling curve; the green solid line represents the fitted line; the green dotted line represents the fitted melting temperature. c. Mean absolute error (MAE) of ΔG, ΔH and Tm between Array Melt data and UV melting data before and after systematic error correction. Gray bars represent errors prior to correction; colored bars represent errors after applying correction with a single offset parameter. d. Direct comparison of measured ΔG, ΔH and Tm between Array Melt and UV melting data. Each datapoint represents one hairpin variant. The gray dots represent measurements prior to correction; the colored dots represent measurements after applying the correction. e. Pearson’s correlation coefficient (*R*) between UV melting data and either Array Melt data or nearest-neighbor model predictions. The colored bars represent the correlation between Array Melt and UV melting data; the black bars represent that between nearest-neighbor model predictions and UV melting data.

### Quantifying nearest-neighbor model performance on non-WC hairpins

After validation of our experimental system, we aimed to examine the performance of the standard nearest-neighbor model on hairpins whose stems are not perfect Watson-Crick pairs. When predicting folding free energies associated with different single mismatch types (**Fig. 3a)**, the nearest-neighbor model achieved an *R*^2^ of 0.61 on the average free energy for each of the types (implying it can explain 61% of the observed experimental variance), or *R*^2^ = 0.31 on all individual variants. However, this model was unable to consistently predict significant variances in folding energies that arise from the same mismatch type with differing flanking sequences (*R*^2^ = −0.61 for T>C mismatches, as shown on the right side panel, *R*^2^ = 0.09 for A>G, *R*^2^ = −0.48 for C>T, and *R*^2^ = −1.66 for G>A) (**Fig. 3b, Fig. S3a**). Indeed, for three out of the four mismatches we analyzed, the nearest-neighbor model introduced extra variance (i.e. negative *R*^2^ value) compared to the class-average prediction, implying that the variance in the nearest-neighbor model’s prediction residual was greater than the actual variance present in the data (**Fig. 3c, Fig. S3b**). While the nearest-neighbor model succeeded in capturing variance for some variant groups with the same mismatch type and nearest-neighbor base pairs, it is unable to capture others (**Fig. S3c**). This limitation of the nearest-neighbor model led us to conclude that there are significant variances in folding energy that are not accounted for by the nearest-neighbor model.

**Figure 3.**
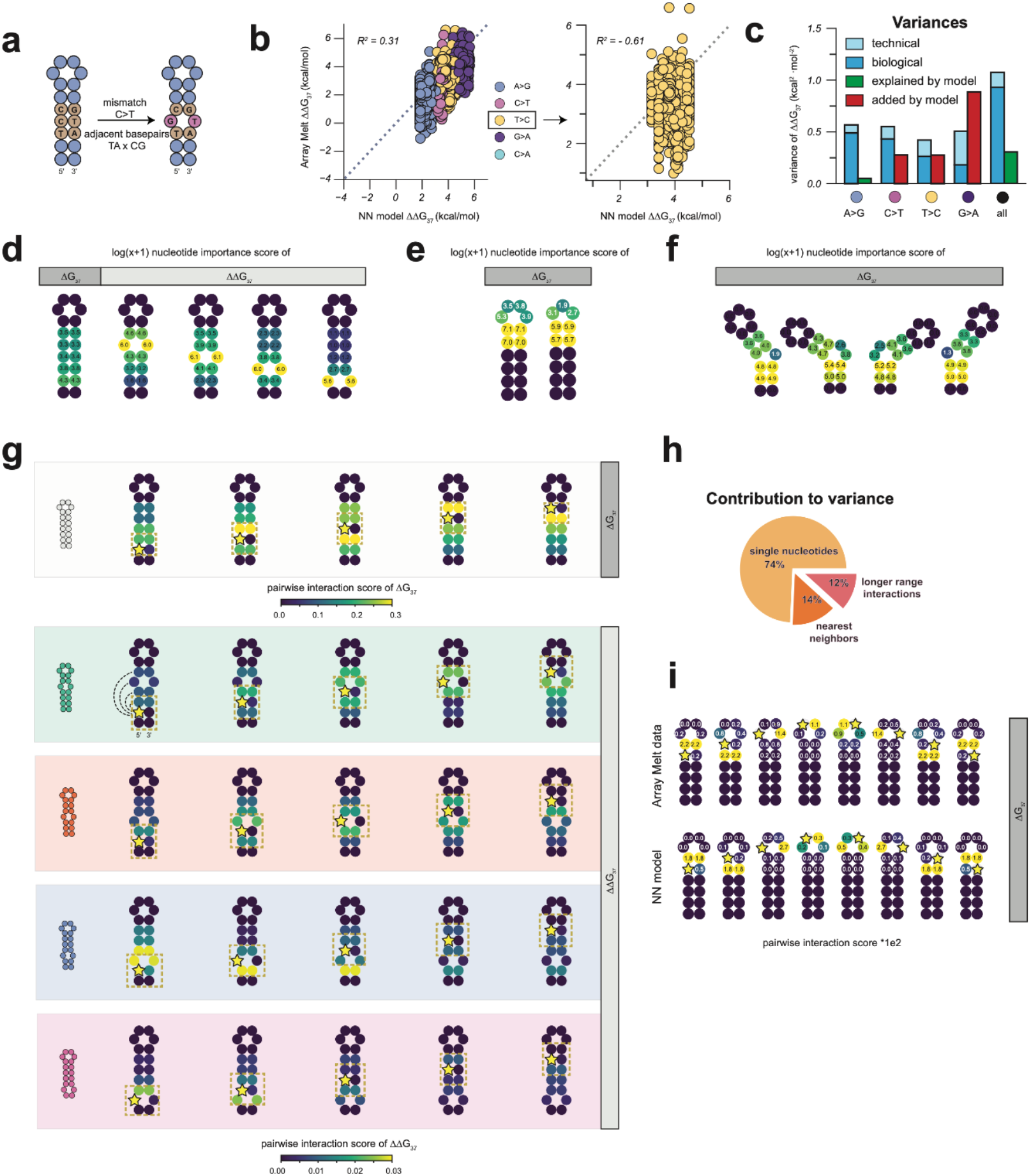
Decomposition of sources of variances in measured energies. a. Schematic representation of mismatch types and ΔΔG calculation for single mismatches. b. Comparison of single mismatch ΔΔG between nearest-neighbor model prediction and Array Melt measurements. Each variant is colored by the identity of the mismatch (left), and the T>C mismatches are zoomed in on the right. c. Unexplained expected biological variance for each group of mismatches and combined. Variance explained or introduced by the nearest-neighbor model are shown in green and red, respectively. d. Nucleotide importance scores for each individual nucleotide in mismatch constructs, grouped by secondary structure. The importance score is calculated as the sum of squares introduced by varying the given nucleotide position. e. Nucleotide importance scores for tetraloops and triloops. f. Nucleotide importance scores for bulges. g. Pairwise interaction scores for each pair of nucleotide locations for Watson-Crick and single mismatch variants. A yellow star indicates the anchor point in each subplot, and the color of each position indicates the strength of the interaction with the anchor. The interaction score is calculated as the R-squared of a linear model given all combinations of the pair as features, subtracted by that of the two individual nucleotides. h. Percentage contribution of single nucleotides, nearest neighbors, and longer-range interactions to the total variance in ΔΔG_37_ for single mismatches. i. Pairwise interaction scores for tetraloops and triloops.

### Influence of sequences on energy across different secondary structures

We hypothesized that some of the observed variance in melting behavior not explained by the nearest-neighbor model may be due to the influence of sequences adjacent to mismatches. To assess this possibility, we calculated the impact of each nucleotide position on folding energy. We first determined a nucleotide importance score at each nucleotide position in different sequence groups with the same target secondary structures by calculating the variance in either ΔG_37_ or ΔΔG_37_ values for the four different identities of the nucleotide at the target position (see **Methods**). While it is possible that variants might adopt alternative secondary structures yet maintain two-state behavior, our analysis still provides valuable insights into how sequences affect folding energy. For single mismatches, we used ΔΔG_37_ values relative to Watson-Crick pairing “parent” variants to regress out the energy expected from Watson-Crick stacks; for other secondary structures, we directly used ΔG_37_ values. As anticipated, nucleotides along the hairpin stem displayed consistent nucleotide importance scores for Watson-Crick pairing variants, and the two mismatching nucleotides have the highest contribution to this folding energy for single mismatch variants, both in the Array Melt data and the nearest-neighbor model (**Fig. 3d, Fig. S3d**). Extending our analysis to hairpin loops and bulges, we observed that specific sequences within these structures critically influence their thermodynamic energy (**Fig. 3e-f**). In triloops and tetraloops, loop center nucleotides had a more pronounced influence on folding energy in the Array Melt data than nearest-neighbor model predictions. We also noted a slight asymmetry between the 5’ and 3’ ends of the loop, regardless of loop size (**Fig. 3e, Fig. S3e**). In bulges, flanking Watson-Crick pairs had greater influence than the bulge sequence itself, with double bulges showing more sequence dependence than singles (**Fig. 3f, Fig. S3f**).

These observations prompted us to examine the energetic interactions between bases by calculating the additional proportion of variance explained in a linear model that considered base interactions (i.e. allowing free parameters for pairwise combinations of bases in the two probed positions) compared to one that treats each position independently (i.e. only allowing free parameters for each individual position). Notably, nucleotides opposite the mismatch showed interactions unaccounted for in the nearest-neighbor model (**Fig. 3g, Fig. S3g**). This interaction score decays with distance along the stem (**Fig. S3h**). Our quantification showed that individual nucleotides, nearest neighbors, and longer-range interactions contribute 74%, 14%, and 12% respectively in this specific case of single mismatches (**Fig. 3h**). In hairpin loops, the interactions between the loop ends and the loop center nucleotides were stronger than expected in tetraloops, but not in triloops (**Fig. 3i, Fig. S3i-j**).

Subsequently, we fit linear regression models to Watson-Crick and single-mismatch variant Array Melt data, varying the length of stack features from one (base pairs only), to two (nearest neighbors) and three (next nearest neighbors). Although prediction errors were similar for nearest- and next-nearest-neighbor models in Watson-Crick variants,next-nearest-neighbor models outperformed the nearest-neighbor ones in single-mismatch variants (**Fig.4a**).

### Revised parameter set for NUPACK integration

We aimed to develop a refined set of parameters to integrate our findings with the NUPACK software^7,20^, thereby enabling more accurate DNA thermodynamic predictions that are easily accessible. We divided Array Melt data into training, validation, and test sets, respectively with 24979, 1315, and 1438 variants, stratified by major variant types as shown in **Fig. 1c**. Initially, we fitted compact linear regression models to the Array Melt data, focusing on structural elements prevalent in our variant library. By allowing the 10 Watson-Crick pairing stack parameters to vary freely (totaling 188 free parameters), we reproduced the established nearest-neighbor parameters set^8^ (**Fig. 4b**). Subsequently, we fixed the values of the 10 Watson-Crick stack parameters and fitted a compact 178-parameter nearest-neighbor model. We compared the single mismatch parameters of our new parameter set with the original *dna04* set and observed similar overall trends. Our new parameters also captured context-dependent variations of GT mismatches and highlighted the instability of CC mismatches, which are shown to be the most thermodynamically unstable DNA single mismatch^21^ (**Fig. 4c, Fig. S4c**). Next, we obtained parameters for tetraloops and triloops by aggregating thermodynamic measurements of relevant Array Melt variants and converting the aggregated ΔΔH and ΔΔG_37_ values to ΔH and ΔG_37_ parameters (**Methods**). We combined these corrected hairpin loop parameters with the compact 178-parameter nearest-neighbor model. This combined model, thereafter referred to as the NUPACK-compatible model, is easily accessible by pointing NUPACK to our parameter set file (**Fig. S4a**; parameter file available in supplementary material). Note that when using NUPACK to predict free energy, we assumed a two-state model in alignment with the Array Melt dataset, which was filtered based on the two-state criterion. Additionally, the ensemble model yielded inferior results compared to the two-state model, both with the original *dna04* and with our new parameter sets (**Fig. S4b**).

**Figure 4.**
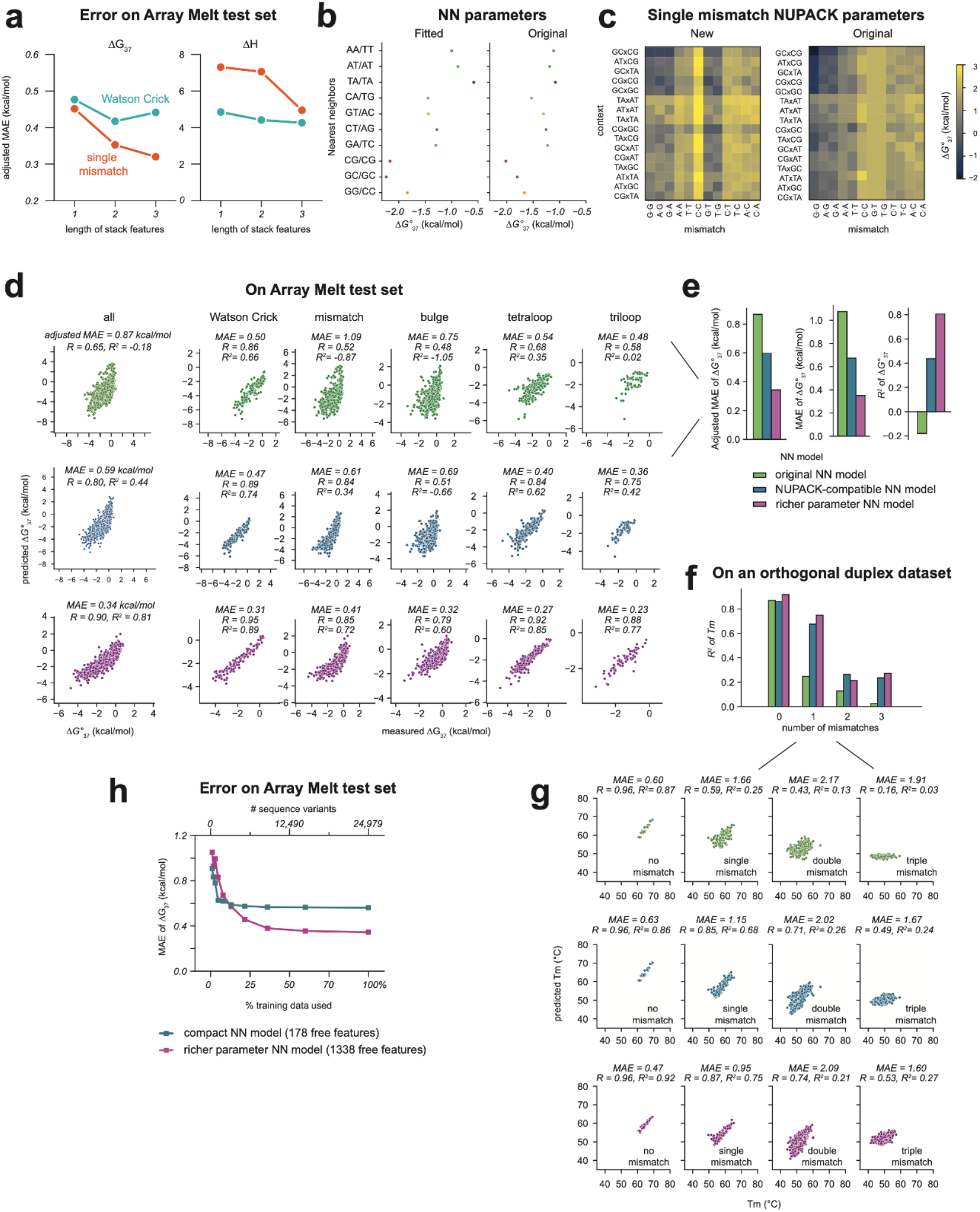
Enhanced linear regression models for DNA folding energetics. a. Adjusted mean absolute error (adjusted MAE) of linear regression models with features based on single base pair, nearest neighbor, or next nearest neighbor stacks, applied to Array Melt test set data. b. Comparison of nearest-neighbor parameters for Watson-Crick pairs fitted from Array Melt data and those reported in the literature^8^. c. Visualization of new and the original *dna04* free energy parameters for single mismatches. “Context” and “mismatch” definitions are consistent with those in **Figure 3a**. d-e. Performance comparison of the original (*dna04*), NUPACK-compatible, and richer parameter nearest-neighbor models on held-out Array Melt data. All calculations were performed at 1M Na^+^ concentration. NN: nearest-neighbor. f-g. Performance comparison of the original (*dna04*), NUPACK-compatible, and richer parameter nearest-neighbor models on an orthogonal dataset of DNA duplexes with varying numbers of mismatches (*Oliveira et al*.^15^). h. Adjusted MAE on Array Melt data plotted as a function of the percentage of training data used to fit the linear regression model.

We evaluated the models on held-out Array Melt data and two orthogonal DNA duplex melting datasets from the literature. To assess model performance, we used both the raw mean absolute error (MAE) and the adjusted MAE, which is more relevant for assessing ΔΔG errors (**Supplementary Notes**). Testing our NUPACK-compatible model on the held-out Array Melt test data resulted in an adjusted MAE of 0.59 kcal/mol for ΔG_37_ (**Fig. 4d-e**). This new parameter set also improved predictions for DNA hairpins designed with random sequences measured with UV melting (**Fig. S4d**). We further asked whether the new parameter set, trained on DNA hairpin data, extends to unseen DNA duplexes. Using NUPACK to predict unseen duplex melting data, we observed a similar performance for Watson-Crick strands with both the original *dna04* and the new parameter set. However, the new parameter set showed improved performance on more complex variants. For a UV melting data set consisting of 384 Watson-Crick pairing DNA duplexes^22^, split evenly into validation and test sets, our parameter set performed comparably to *dna04* (**Fig. S4e**). On a dataset from *Oliveira et al*. containing DNA duplex melting temperatures^15^ with varying numbers of mismatches (also split evenly between validation and test sets), both parameter sets had a similar adjusted MAE of around 0.6 °C for Watson-Crick duplexes on the test set; while for single mismatches, the new parameter set achieved a much lower adjusted MAE of 1.15 °C (*R*^2^ = 0.68) compared to 1.66 °C (*R*^2^ = 0.25) with *dna04*. The improvement was even greater for double and triple mismatches (**Fig. 4f-g**). In conclusion, this updated parameter set, while simple, enhances the prediction accuracy of NUPACK for DNA secondary structures, particularly those with mismatches and hairpin loops. This improvement extends to orthogonal DNA duplex data that the model was not trained on.

### Enhanced linear model with rich parameterization

Building on this NUPACK-compatible model, we next developed an enhanced linear model that incorporates a richer parameter set (“richer parameter model”), allowing still more accurate and comprehensive prediction of DNA thermodynamics on unseen validation data. Because this model has more parameters than are implemented in the NUPACK, it is not immediately compatible with the software. We fit this model with 1338 free parameters, incorporating still more context information (including interactions between the two stacks flanking a mismatch, bulge sequence identity, and full hairpin loop sequence for triloops and tetraloops, see **Methods**), to the Array Melt training data. This rich parameter nearest-neighbor model yields an even better performance both on the Array Melt test data (adjusted *MAE* = 0.34 *kcal*/*mol, R*^2^ = 0.81) and the external duplex datasets (**Fig. 4d-g, Fig. S4e**). Notably, when comparing the prediction error on Array Melt validation dataset as we titrate the percentage of training data used to fit the model, the 178 parameter linear regression model used to build the NUPACK-compatible model plateaus with just a small fraction of training data used, while the 1338 parameter model requires more measurements to plateau, but still does so after being fit on less than half of the training data set (**Fig. 4h**). Given the relatively overdetermined nature of even this rich parameter nearest-neighbor model, this behavior is expected, and clearly demonstrates that increasing data size allows for more variance to be captured by a more expressive model.

### Graph neural network (GNN) model for DNA energy prediction

Because our rich linear model likely had insufficient power to capture the biological variance in our data, we developed a graph neural network model^23,24^ specifically designed for predicting DNA energy. In this model, a DNA molecule is represented as a graph where nodes represent nucleotides and edges represent chemical bonds. Nucleotides (A, T, C, or G) and chemical bonds (either 5’ to 3’ backbone, 3’ to 5’ backbone, or hydrogen bond) are one-hot encoded as node and edge features, respectively. Taking these graph representations as inputs, the GNN iteratively refines node embeddings across graph convolutional layers. Essentially, nodes exchange information through different edge types, allowing, over multiple layers, for the embedding of one node to encapsulate information from the entire graph. Investigation of various types of graph convolutional layers revealed that the graph transformer layers^25^ outperformed other architectures, such as graph attention layers^26^ (**Fig. S5a**). After graph convolution, each node possesses a learned embedded feature, reflecting not only the original node features, but also the edges and overarching graph structures. To embed this graph into a fixed-length vector representation, we employed a global pooling layer employing the Set2Set algorithm^27^, which used a Long Short-Term Memory (LSTM) processing block. This block systematically ingests node embeddings while updating its internal state, resulting in a vector representation of the entire graph that is order-invariant to the input node features. This processed summary vector is then fed into a small, fully connected neural network, which regresses for ΔH and Tm values (**Fig. 5a**).

**Figure 5.**
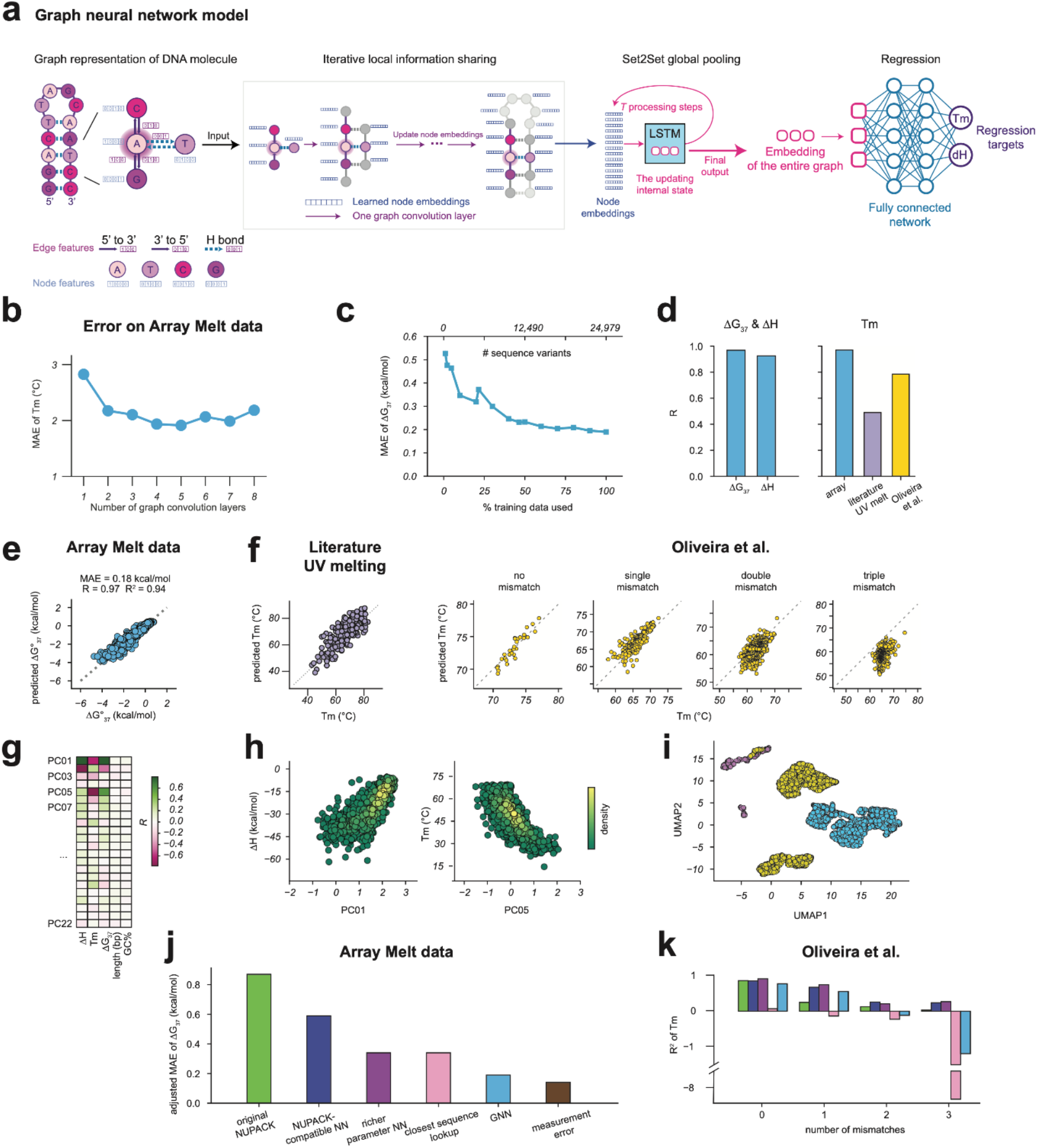
Graph neural network model for DNA folding thermodynamics. a. Schematic representation of the graph neural network (GNN) architecture. b. Mean absolute error (MAE) of melting temperature on Array Melt data as a function of the number of graph convolution layers in the GNN. c. MAE of ΔG_37_ on Array Melt data as a function of the percentage of training data used to fit the GNN. d. Pearson’s *R* correlation coefficient between GNN predictions and held-out Array Melt data, literature UV melting data, or Oliveira et al. duplex melting data. e. Scatter plot comparing GNN predictions with held-out Array Melt data. Each dot represents a single sequence variant. f. Scatter plot comparing GNN predictions with literature UV melting data or Oliveira et al. duplex melting data. g. Pearson’s *R* between principal components of the learned variant embeddings and measured thermodynamic parameters or variant properties. h. Comparison between the most correlated principal components and ΔH or Tm. i. UMAP visualization of learned variant embeddings after aggregation, colored by dataset (Array Melt, literature UV melting, or Oliveira et al. duplex melting data). j. Benchmark of all models on held-out Array Melt data, compared to the measurement error of Array Melt. k. Benchmark of all models on the orthogonal Oliveira et al. dataset, plotted by the number of mismatches. Colors correspond to models as in **Fig. 5j**.

We trained this end-to-end model on Array Melt data. A portion of the Array Melt data was held out as validation and test set. We first explored the effect of the number of graph convolutional layers on prediction accuracy (**Fig. 5b, Fig. S5b**). Performance converged at around 4 graph convolutional layers, and did not improve beyond 4 layers, consistent with our observations in **Fig. S3h** that pairwise nucleotide interactions span around a distance of 4 nt. When evaluating model performance over an increasing percentage of training data used as in **Fig. 4h**, we observed a slower plateauing of the curve, indicating that the model can make use of this large-scale data set (**Fig. 5c**). The trained GNN achieved prediction error of 0.18 kcal/mol in *dG*_37_ (*R* = 0.97, *R*^2^ = 0.94), compared to 0.34 kcal/mol of a baseline model that simply looks up the k=8 closest variants in the training set and gives a weighted average of the lookup values as prediction (**Fig. 5d-e, Fig. S5c-d**), and the GNN predictions are consistently high quality for different types of sequence variants (**Fig. S5e**). Interestingly, although the GNN was trained entirely on graph representations of DNA hairpins without knowing the nature of backbones and base pairing, it is able to extend to the annealing and melting properties of DNA duplexes (**Fig. 5f**). While the GNN performance is slightly worse than our rich parameter nearest-neighbor model (**Fig. 4f-g, Fig. S4e**) on DNA duplexes, this level of performance suggests that the neural network has grasped the “gist” of the physical system of DNA folding thermodynamics and can extrapolate to non-hairping annealing contexts. Compared to the slow decrease of prediction error on the Array Melt data, prediction error rapidly decreased within around the first 20 epochs and then plateaued for the two DNA duplex datasets (**Fig. S5f**), likely because of the relatively low complexity of those datasets.

To better understand how the GNN makes predictions, we analyzed the intermediate activation from the output of the Set2Set aggregation layer. The GNN projects each DNA molecular graph into a 250-dimensional embedding vector. Many of the top principal components of the embeddings correlate strongly with predicted thermodynamics parameters (*R* = 0.8 between PC1 and ΔH, *R* = −0.77 between PC5 and Tm, and *R* = 0.79 between PC1 and ΔG_37_), but none with other qualities of the sequence variants including DNA length or GC content (**Fig. 5g-h**), underscoring the GNN’s focus on thermodynamics. When examining embeddings from different datasets (Array Melt, literature UV, and Oliveira et al.), each dataset’s distinct design structure mapped to unique areas in a two-dimensional UMAP visualization (**Fig. 5i**). This observation indicates that the embeddings capture more than just predicted thermodynamics like melting temperatures (**Fig. S5g**), and that the learned embedding space encompasses sequence and structure details beyond mere parameter prediction.

Finally, we compared prediction errors across various models. The graph neural network is the top performer on Array Melt test data, while the 1778-parameter richer parameterization nearest-neighbor model performs best when applied to duplex data. Precision of the GNN is noteworthy, nearing the uncertainty range of bootstrapped experimental measurements (mean of uncertainty is 0.14 kcal/mol in Δ*G*_37_), highlighting its effectiveness in capturing complex relationships in DNA sequences and structures (**Fig. 5j, Table 1**).

**Table 1.** Model benchmarks. MAE: mean absolute error. Units of adjusted MAE and MAE are kcal/mol for ΔH and Δ_37_ and °C for Tm. Numbers in the “Measurement” column are means of bootstrapped uncertainties from measurement.

|  |  | Model | Original<br>NUPACK | NUPACK-compatible | Rich<br>parameterization | Closest<br>lookup | Graph neural<br>network | Measurement |
| --- | --- | --- | --- | --- | --- | --- | --- | --- |
| Array | $\Delta H$ | Adjusted MAE | 8.59 | 6.27 | 4.97 | 5.27 | 3.13 | 3.09 |
|  |  | MAE | 14.33 | 10.79 | 5.03 | 4.03 | 3.13 | 3.09 |
| | | $R^2$ | 0.12 | 0.48 | 0.64 | 0.77 | 0.86 | NA |
| | $T_m$ | Adjusted MAE | 7.94 | 6.52 | 5.09 | 3.72 | 1.80 | 1.28 |
|  |  | MAE | 8.38 | 6.53 | 5.65 | 3.72 | 1.80 | 1.28 |
| | | $R^2$ | -0.06 | 0.26 | 0.20 | 0.78 | 0.94 | NA |
| | $\Delta G_{37}$ | Adjusted MAE | 0.87 | 0.59 | 0.34 | 0.34 | 0.19 | 0.14 |
|  |  | MAE | 1.07 | 0.67 | 0.35 | 0.34 | 0.18 | 0.14 |
| | | $R^2$ | -0.15 | 0.44 | 0.81 | 0.81 | 0.94 | NA |
| Oliveira<br>et.al | $T_m$ (all) | Adjusted MAE | 2.06 | 1.77 | 2.76 | 5.12 | 5.30 | NA |
|  |  | MAE | 5.43 | 4.70 | 10.20 | 9.01 | 5.56 | NA |
| | | $R$ | 0.81 | 0.86 | 0.66 | 0.35 | 0.78 | NA |
| | | $R^2$ | 0.65 | 0.73 | 0.42 | -1.06 | 0.49 | NA |
| | $R^2$<br>of $T_m$ | No mismatch | 0.87 | 0.86 | 0.92 | 0.07 | 0.78 | NA |
|  |  | Single | 0.25 | 0.68 | 0.75 | -0.14 | 0.56 | NA |
|  |  | Double | 0.13 | 0.26 | 0.21 | -0.23 | -0.12 | NA |
|  |  | Triple | 0.03 | 0.24 | 0.27 | -8.33 | -1.21 | NA |

## Discussion

In this work, we developed an accurate, scalable fluorescence-based method for measuring the melting of DNA hairpins (Array Melt) and used it to generate the largest-to-date DNA folding thermodynamics dataset. The Array Melt method, with its high throughput and precise measurement capabilities, allows for a more detailed exploration of the thermodynamics of diverse DNA structures, including those with mismatches, bulges, and various loop motifs. The scale of this data set revealed the significant role played by sequences adjacent to mismatches and within loops to the thermodynamic stability of hairpins. This observation implies that traditional models that parameterize only the immediate nearest neighbors must be underpowered to capture the variance observed in these molecular variants. To this end, we developed three quantitative folding energy models of increasing complexity, including revised parameterization of the linear regression model compatible with NUPACK, a richer nearest-neighbor model especially accurate for duplex annealing of mismatched molecules, and graph neural network (GNN) model that accells at predicting the melting behaviors of hairpins. To our knowledge, this GNN model is the first deep learning model applied to DNA folding. These results support the notion that while traditional nearest-neighbor models are accurate in predicting Watson-Crick stack behaviors, they fall short in accurately modeling the thermodynamics of more complex DNA structures, such as hairpins and interior loops. We anticipate that future library designs will be more oriented towards machine learning, enabling a more comprehensive and less biased sampling of the DNA sequencing space.

Overall, by providing a more accurate model for DNA folding thermodynamics, we pave the way for improved design of biotechnological tools and applications. For example, our models will provide a more accurate starting point for *in silico* design in areas such as qPCR primer design, DNA oligo hybridization probe design, and DNA origami. However, the study’s focus on two-state folding presents a notable limitation, although our comparison between the two-state and ensemble methods (**Fig. S4b**) revealed greater accuracy of the former in predicting Array Melt measured thermodynamic parameters. The inferior performance of the ensemble method may be attributed to the inference methods of parameters. Additionally, the experimental design of the datasets used to derive nearest-neighbor models may also play a role. Therefore, interpreting nearest-neighbor model predictions and dynamic programming algorithm outputs as “true” energies associated with monolithic secondary structures warrants careful consideration. In this study, the two-state folding energies are better interpreted as an aggregated average across multiple folding states, rather than that of a single folded microstate within this ensemble. We anticipate that future analysis incorporating experimental design, data preprocessing, and parameter inference^28^ techniques that consider ensemble folding could result in further improvements in prediction accuracy. This is particularly relevant when predicting probabilities of microstates of dynamic molecules is important.

This work also indicates that cloud labs can provide researchers with practical access to specialized instruments and support, addressing traditional challenges in equipment access and maintenance. These platforms also facilitate streamlined data collection and metadata tracking, improving open data and experiment reproducibility. This study represents an early instance of using commercial cloud labs in research setting^29^, suggesting potential broader academic use.

Finally, while our measurements were carried out at a single monovalent salt concentration, large-scale measurements exploring the stability of DNA structure to the concentration and type of monovalent or divalent cation would be relatively straightforward. Furthermore, the potential application of our method to RNA structures presents an exciting avenue for future research. Given the diverse and critical biological functions of RNA, which are heavily dependent on its secondary structure, applying our high-throughput and precise thermodynamic measurement technique could unveil new dimensions in RNA biology and facilitate the development of RNA-based therapeutics.

## Acknowledgement

We thank Dr. Rhiju Das (Stanford University) for comments on the manuscript; Dr. Mark Schnitzer (Stanford University) for supporting the project; Dr. Niles Pierce and Dr. Mark Fornace (Caltech) for answering NUPACK-related questions. We thank Emerald Cloud Lab (ECL) for the UV melting experiments, with special attention to Malav Desai and Ben Kline for tutorials and technical support of Symbolic Lab Language (SLL) programming and Brian Frezza for funding.

This work was supported in part by NIH grants R01GM111990, P50HG007735, R01HG009909, P01GM066275, UM1HG009436 and R01GM121487 to W.J.G. W.J.G. acknowledges support as a Chan Zuckerberg Investigator. E.M. acknowledges support from the Swedish Research Council (grant 2020-06459), the Foundation Blanceflor, and the Science for Life Laboratory (SciLifeLab).

## Author contributions

Conceptualization, W.J.G., Y.K., E.S., and H.K.W.; Methodology, Y.K, E.S.; Software, Y.K. and H.K.W.; Validation, Y.K; Formal Analysis, Y.K.; Investigation, Y.K, E.S., H.K.W., W.R.B., and A.H.; Data Curation, Y.K.; Writing - Original Draft, Y.K. and W.J.G.; Writing - Review & Editing, Y.K., E.S., H.K.W., E.M., and W.J.G.; Visualization, Y.K.; Supervision, E.M. and W.J.G.; Funding Acquisition, W.J.G.

## Methods

### Library assembly and sequencing

Designed library variants were synthesized into DNA by Twist Biosciences (South San Francisco, CA). Sequence map of the library is provided in supplementary information as *array_melt_library*.*gb*, where the “filler sequence” is a 5’ partial segment of AACAACAACAACATACTAACAACAACATAACAAATCAAAA, its length and the variable region adding up to 40 bp, such that all synthesized fragments have the same length despite different variable region length. Reverse complement of the sequence map was ordered as a synthesized oligo pool, amplified using internal primers to enrich for full-length library variants. The PCR reaction consisted of 1 in 8 dilution of the synthesized oligo pool (final concentration 10 nM, but concentration of full-length library variants could be much lower), 200 nM of each primer (T7A1library, D-TruSeqR2 Table S1), 1x Phire Hot Start II PCR Master Mix (ThermoFisher Scientific F125L). The reaction proceeded for 9 cycles of 98°C for 10 seconds, 56°C for 30 seconds, and 72°C for 30 seconds. Reaction mixtures were purified using QIAquick PCR Purification Kit (Qiagen 28104) to remove primers and proteins, then eluted into 20 uL elution buffer.

After initial amplification, the library was amplified with primers to bring in sequences compatible with Illumina sequencing. This five-piece assembly PCR included two outside primers and two adapter sequences. The PCR reaction consisted of 1 μl of the previous reaction, 137 nM of outside primers (*short_C* and *short_D*; Table S1), 3.84 nM of the adapter sequences (*C-i7pr-bc-T7A1* and *D_TruSeqR2*; Table S1), 1x Phire Hot Start II PCR Master Mix (ThermoFisher Scientific F125L). The reaction proceeded for 14 cycles of 98°C for 10 seconds, 56°C for 30 seconds, and 72°C for 30 seconds. Reactions were purified using the QIAquick PCR Purification Kit and quantified with a Qubit Fluorometer (ThermoFisher Scientific).

Sequencing was run with MiSeq Reagent Kit v3, 150 Cycles (MS-102-3001). The amplified library was diluted and pooled to 1.16 nM final concentration in the pool (27% sample), together with a final concentration of 2.8 nM PhiX control v3 (70% sample, Illumina), 0.04 nM fiducial marker (1% sample), 0.04 nM Cy3 no RNA control (1% sample), and 0.04 nM single-strand fluorescence control sequences (1% sample, not included in analysis), adding up to a total sample concentration of 4 nM. Paired-end sequencing was done with a custom read 1 primer (*stall_R1_primer*) and a standard read 2 primer to cover the variable region in both read 1 and read 2 directions.

### Imaging station setup

An imaging station was used to image the Miseq chip at increasing temperatures. This station was built from a combination of custom-designed parts from a disassembled Illumina genome analyzer IIx^18,30^. Two channels were employed: the “red” channel used the 660 nm laser and 664 nm long pass filter (Semrock BLP01-664R-25), and the “green” channel used the 532 nm laser and 590 nm band pass filter (Semrock FF01-590/104-25). All images were taken with 600 ms exposure times at 150 mW fiber input laser power. Focusing at each temperature was achieved by sequentially adjusting the z-position and re-imaging the four corners of the flow cell; the adjusted z-_positions_ were then fit to a plane. We chose Cy3 as the fluorophore because the “green” channel is relatively more stable in our instrument setup.

Post-sequencing, the chip was washed with Cleavage buffer (100 mM Tris-HCl, 125 mM NaCl, 0.05% Tween20, 100 mM TCEP, pH 7.4) to remove residual fluorescence from the reversible terminators used in the sequencing reaction at 60°C for 5 minutes. Any strands of DNA not covalently attached to the surface of the chip was removed by washing in 100% formamide at 55°C. The resulting single-stranded DNA fragments were incubated with 500 nM of the oligo *Biotin_D_Read2* and *red_oligo* (Table S1) in Hybridization buffer (5x SSC buffer (ThermoFisher 15557036), 5 mM EDTA, 0.05% Tween20) for 15 minutes at 60 °C, subsequently the temperature was lowered to 40 °C for another 10 minutes. The *red_oligo* is conjugated to Alexa647 and binds a sparse subset of “fiducial” sequences on the chip, whose fluorescence signal is used for initial image registration to the sequenced tiles.

### Measuring melt curves on the chip

The Cy3 labeled *fluor_oligo* was annealed at 40°C for 5 minutes, then the Black Hole Quencher (BHQ)-labeled oligo *quench_oligo* (**Table S1**) using the same protocol. The chip was imaged after each step to ensure that hybridization had occurred through increase or quenching of signal in the corresponding channel. The chip was then rinsed with melt buffer (50 mM Na-HEPES pH 8.0, 25 mM NaCl, total Na^+^ concentration about 88 mM at pH 8.0). We chose HEPES buffer because it has one of the lowest sensitivities to temperature (-0.014 ΔpK_a_/°C) among the most commonly used buffers in biological experiments^31^.

To quantify the melt curves, image station temperature was then lowered to 20°C. For each temperature point, the system was allowed to equilibrate to the new temperature (approximately 5 minutes) before refocusing and imaging. The temperature was raised in 2.5°C increments to a maximum temperature of 60°C. Detailed protocol can be found in XML imaging scripts in the supplementary information.

### Processing sequencing data

Sequencing data from Illumina Miseq was processed to extract tile and coordinates of each sequenced cluster. Forward and reverse paired-end reads were aligned using FLASH^32^ with default settings. Consensus sequences from FLASH were aligned to the reverse complement sequences of *fluor_oligo* and *quench_oligo* using a Needleman-Wunsch alignment (nwalign3). For consensus sequences that successfully aligned to both with p-value <1e-3, evaluated as described in^33^, the variable region was extracted as the region between the two flanking regions and aligned to sequences in the reference library. A reference sequence was assigned based on the best-scoring alignment with a p-value < 1e-6. The resulting clusters were used for fluorescence quantification in the downstream data analysis.

### Fluorescence data processing and image fitting

Fluorescent melt curve images were mapped to sequencing data from the Illumina Miseq. First, sequencing data was processed to extract the tile and coordinates of each sequenced cluster. The coordinates were used to generate synthetic images of clusters, which were registered to the fluorescent images iteratively to map the coordinates to the images at sub-pixel resolution^34^. Once the identities of the clusters were determined, each cluster was fitted to a 2D normal distribution to quantify its fluorescence^30^.

### Distance sensitivity curve

The repeat controls used to plot the distance sensitivity curve in Fig. 1d are repeats of varied lengths of the following sequences: poly-A, T, AT, AAC, AAG, AAT, AC, ACC, AG, AGG, TC, TG, TTA, TTC, and TTG.

The theoretical curve fitted to the distance-to-fluorescence curve in Fig. 1d is 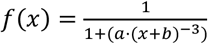, where x is the length in nt, *f*(x) is the dimensionless fluorescence at length x, normalized to maximum fluorescence, *b* is a fitted parameter in nt, which accounts for systemic distance offset, and *a* is another fitted parameter:

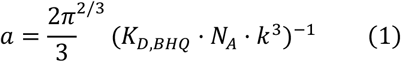

where *K*_*D,BHQ*_ is the dissociation constant for the fluor-quench pair, *N*_*A*_ is Avogadro’s number, and *k* = 0.64 × 10^−9^ *m* ⋅ *nt*^−1^ is a distance conversion factor from nt to physical distance for ssDNA. The fitted parameter values are *a* = 71.19, *b* = 0.08, and *K*_*D,BHQ*_ = 145.6 *mol* ⋅ *L*^−1^. Using Δ*G* = *RT* ⋅*ln K*_*D,BHQ*_, we have *dG*_37_ = 3 *kcal*/*mol*.

### Fluorescence normalization

#### Size normalization

To reduce inter-cluster variation in fluorescence measurements, we normalized the Cy3 fluorescence during the melt curve measurements by the total amount of ssDNA in that cluster, measured as Cy3 fluorescence right after hybridization of *fluor_oligo* but before hybridization of *quench_oligo*. The normalizing factor Cy3 signal was clipped to the 1st and 99th percentile of the total distribution.

#### Temperature dependence and photobleaching normalization

Across the timespan of the experiment, fluorescent signals are dependent on temperature and undergo photobleaching. To correct for these factors, we further normalized the size-normalized signals of clusters to median signals of control variants designed to have constant distances between the Cy3 fluorophore and the quench, including 1) “super stable stem” variants with long GC stems that are not supposed to melt within the experimental temperature range and show minimum fluorescence signals, and 2) “long repeat” controls that are at least 39 nt long, not supposed to form a hairpin, and show maximum fluorescence signals. For both control variant groups, only the variants whose median signal across clusters fall within the 5th and 95th percentile of all median signals at each temperature point within that control variant group are used. Each variant is allowed to fall outside the percentile range at 1 temperature point at most. This step filters out the variants not exhibiting the expected behavior. Afterwards, the median values at each temperature point of both groups are taken and used to normalize the entire dataset. **(Fig. S1d)**

#### Direct melt curve fitting

The melt curve fitting pipeline was adapted from^34^. Briefly, the direct fitting process was carried out in the following steps: 1) Directly fit normalized signals of single clusters; 2) Estimate the distribution of *f*_*max*_ and *f*_*min*_ from single cluster fit results; 3) Refine the fit at the variant level with bootstrapping. The details are described below. The fitting process is available as a snakemake^35^ pipeline at https://github.com/keyuxi/array_analysis.

##### 1) Single cluster fitting

**Equation (2)** was directly fitted to normalized signals of single clusters with minimal constraints, where *T* and *T*_*m*_ are temperatures in Kelvin, and *k*_*B*_ = 0.0019872 is the Boltzmann constant. For initial values, Tm was set to the temperature point where the normalized signal is closest to 0.5, ΔH to -40, *f*_*max*_ to 1, and *f*_*min*_ to 0. The python package *lmfit* (v1.0.3)^36^ was used for least-squares fitting.

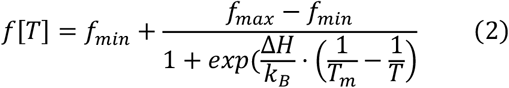

##### 2) Estimating distributions of f_max_ and f_min_

We used the following empirically gated criteria to decide if a single cluster fit was successful: *R*^2^ of the fit at least 0.5, fitted *f*_*min*_ between -1 and 2, the estimated standard error of the best-fitted parameter value *σ*_*fmin*_ less than *f*_*min*_ + 1, fitted *f*_*max*_ between 0 and 3, *σ*_*max*_ less than *f*_*max*_,*σ*_*Tm*_ less than 10 K, and *σ*_Δ*H*_ less than 100 kcal/mol. A p value is calculated for each variant with a null hypothesis of 25% successful single cluster fits as in^34^. The single clusters are then aggregated to the variant level. Median values of the parameters of each variant were calculated. To select for variants used for estimation of *f*_*max*_ the previously mentioned single cluster criteria were combined with the following extra criteria: 1) be a good fitter, defined as having a success p value less than 0.01; 2) be unfolded at high temperatures, defined as having a fitted *f*_*max*_ value of at least 0.5, and 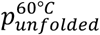, the portion of unfolded molecules in the cluster calculated from the fitted parameters, of at least 0.975. Similarly, for *f*_*min*_ we select for variants with a p value less than 0.01, *f*_*min*_ less than 0.05, and 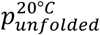 at most 0.025. We then fitted cluster-per-variant-dependent Gamma distributions to *f*_*max*_ and *f*_*min*_ as in^34^.

##### 3) Fit refinement

For each variant, we separately decided whether to enforce *f*_*max*_ and *f*_*min*_ distributions to correct for variants that never become fully unfolded or folded during the experiment. Specifically, we calculated the lower bound for *f*_*max*_ as

*μ*_*fmax*_ − 10 ⋅ *σ*_*fmax*_, where *μ*_*fmax*_ and *σ*_*fmax*_ are the mean and standard deviation calculated from the Gamma distribution with the according number of clusters per variant from the previous step, and the upper bound for *f*_*min*_ as *μ*_*fmin*_ + *σ*_*fmin*_. Here, we set the margin for *f*_*max*_ larger to enforce global *f*_*max*_ less, as *f*_*max*_ is more variable between variants than *f*_*min*_ because of sequence-dependent effects. If the median signal at the highest temperature point is less than the lower bound for *f*_*max*_, we enforce *f*_*max*_; likewise, if the median signal at the lowest temperature point is greater than the upper bound for *f*_*min*_, we enforce *f*_*min*_.

We fitted each variant by bootstrapping the clusters within that variant. At each resampling step, we draw a *f*_*max*_ or *f*_*min*_ value from the global distribution if enforced or keep *f*_*max*_ or *f*_*min*_ floating otherwise, and fit **equation (2)** to the median signal of the drawn clusters, using the single cluster fit results as the initial parameters. Then, we calculate Δ*G*_37_ and ΔS for each resampling step:

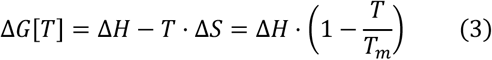

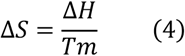

where *T* is temperature in Kelvin. We repeated the process 100 times and obtained a 95% confidence interval and standard deviation for each parameter, ΔH, Tm, ΔG_37_ and ΔS.

#### ΔG Line fitting

We also fitted the data with an alternative line fitting to obtain heuristics for two-state behavior. We first re-normalize the signal *f*[*T*], previously normalized to initial fluorescence after hybridization as aforementioned, with fitted *f*_*max*_ and *f*_*min*_. We clip *f*_*norm*_[*T*] = (*f*[*T*] − *f*_*min*_)/(*f*_*max*_ − *f*_*min*_) between 0 and 1 with an *ϵ* = 0.01 to avoid problems in the next step. We transformed *f*_*norm*_ [*T*] to Δ*G*[*T*] with

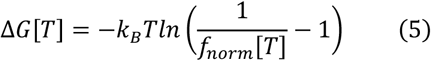

Error propagation was calculated with the uncertainties python library. Finally, we fitted with a line with free parameters ΔH and Tm:

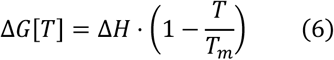

If the variant exhibits two-state behavior, Δ*G*[*T*] should fit well to a linear line. We first fit with RANSAC^37^ to identify outliers, then re-fit the inliers with ordinary least squares linear regression for the final parameters.

#### Filtering and combining data replicates

Each variant measured and fitted to curves in a replicate experiment was first filtered by the criteria: at least measured in 5 clusters, bootstrapped error in ΔG_37_ is less than 2 kcal/mol, error in Tm less than 25°C, error in ΔH less than 25 kcal/mol, and curve fitting RMSE less than 0.5. They were then compared to the line fitting results. To be labeled as passing two-state criteria, the fitted ΔH value of a variant between the 2 methods need to agree within 20%, Tm within 2°C, the reduced *X*^2^ of the line fit below 1.5, and the number of inlier data points in line fitting greater than 10. The replicates were then combined as described in^34^. Briefly, for each parameter (ΔH, Tm, Δ*G*_37_, or ΔS), its measurements *X*_*i*_ and bootstrapped error *σ*_*i*_ in each replicate, the combined error is 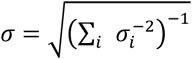 and the combined parameter 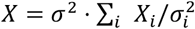, where the sum is over the replicates whose data are not missing for the given variant. The missing values were ignored. We then apply a second filter for two-state behavior. A given variant needs to be labeled as passing two-state criteria in at least 2 replicates to be kept. We combined 4 such replicate experiments, including 3 melting and 1 cooling curve measurements from 2 different chips (see table for detail). We apply a final filter on the combined variants to only keep those whose Tm are between 25 and 55°C and fall within the dynamic range of the system.

### UV melting

#### Designing sequences for UV melting and quality control

We randomly selected hairpin sequences from each category of the Array Melt library for validations in **Fig. 2** and designed random 16-mers that would fold into a hairpin with the secondary structure ((((((….)))))) for **Fig. S4d**, filtering those with misfold secondary structures with NUPACK. DNA oligo samples were synthesized by Integrated DNA Technologies (IDT) and were run through Ion Exchange High Pressure Liquid Chromatography (IE-HPLC) remotely on Emerald Cloud Lab (ECL) with DNAPac PA200, 9x250 mm Semi-Prep column (Thermo Fisher Scientific 063421), and the fractions were measured by a UV-Vis detector at 260 nm. Only the oligos with a single peak were used for data analysis.

#### Running cloud lab UV melting experiments

UV melting experiments were run remotely on ECL. The Manifold data objects could be found at the data object links in supplementary table 2. DNA oligos ordered from IDT were shipped to ECL and resuspended in high concentration stock solutions. For hairpins, the DNA stock solutions are first snap freezed right before each experiment, being placed at 90°C for 5 min and moved to 4°C for 30 min before diluted into working concentrations; for duplexes, the DNA stock solutions are first diluted into working concentrations, then placed on 90°C for 5min and left annealing for 30 min to 1 hour. Melt curves are measured at varied concentrations, where the range is 3 to 12 uM for hairpins and 72 to 96 uM for duplexes. Hairpin data used for method validation in **Fig. 2** and model comparison in **Fig. S4d** were all measured at 6 uM or 9 uM. Samples were measured on a Cary 3500 UV/Vis spectrophotometer, using micro scale black walled UV quartz cuvettes with a pathlength of 10 mm. Samples were measured for 2 to 3 melt-cooling cycles with a temperature ramp rate of 0.8°C and 15 min equilibrium time between each melt and cool sweep. Temperatures were measured with an immersion probe. Data points were acquired every 1 °C at 260 nm from 5°C to 85 or 95 °C.

#### Analyzing UV melting curves

The UV melting curves were fitted to a modified version of **equation (2)** with an extra term to account for baseline slope:

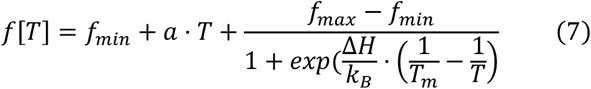

where f[T] is the absorbance, *a* is a free parameter for the slope, and the rest the same as in **equation (2)**. For each individual melt or cooling curve, we first found the temperature where the derivative 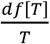 is maximum, that is, 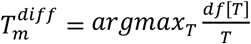, as the initial value of *T*_*m*_. We fixed the value of *T*_*m*_ when fitting **equation (7)** in the first round and let the parameters *a, f*_*max*_, *f*_*min*_, and Δ*H* float. Afterwards, we use the best-fitted values from the previous round as the initial values of the parameters and allow *T*_*m*_ to float in the second round of fitting. This two-step fitting process empirically improved the fitting RMSE. We selected the curves with an RMSE less than 0.015, *σ*_Δ*H*_ < 10 And 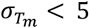 and aggregated the fitted Δ*H* and *Tm* values of the melt and cooling curves by each sample tube with 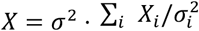 and 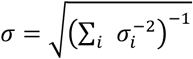 as for the array experiments. We then calculated Δ*G*_37_ and Δ*S* from the aggregated Δ*H* and *T*_*m*_ using equations (3) and (4), and used the uncertainty propagation equations *σ*_Δ*G*_ = −Δ*S* ⋅ 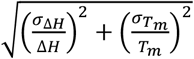 and 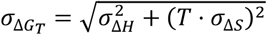 to calculate the measurement uncertainties.

#### Salt correction

All salt corrections used the equation from^38^:

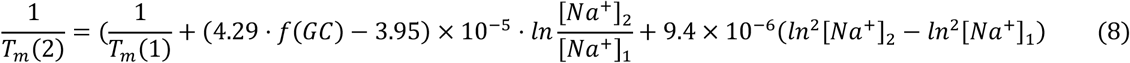

where *T*_*m*_ (1) and *T*_*m*_ (2) are the melting temperatures before and after salt correction in Kelvin, [*Na*^+^]_1_ and [*Na*^+^]_2_ the corresponding sodium concentrations, and *f*(*GC*) the fraction of GC in the sequence.

#### Variance analysis

##### Mismatch ΔΔG_37_ calculation

For each mismatch, we find its parent sequence variant, which is a hairpin with Watson-Crick pairing stem and can derive into the mismatch by mutating a single nucleotide. The mismatches are categorized by the identities of those nucleotide substitutions. We then calculated ΔΔ*G*_37_ of mismatches by subtracting Δ*G*_37_ of the parent variant from that of the mismatch variant. C>G single mismatches were excluded from the variance calculations because of insufficient number of data points (only 2 variants).

##### Expected biological variance and model explained or introduced variance

For a dataset of *n* sequence variants with *n* corresponding *d*Δ values *y*_1_, …, *y*_*n*_, measurement uncertainty values *σ*_1_, …, *σ*_*n*_, and nearest-neighbor model prediction values *ŷ*_1_, …, *ŷ*_*n*_, we have total variance 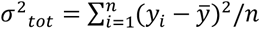 and technical variance 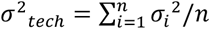. The expected biological variance *σ*^2^_*bio*_ = *σ*^2^_*tot*_ – *σ*^2^_*tech*_. Model variance is the variance of prediction values 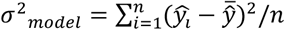, and residual is 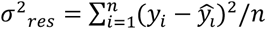. From these we calculate variance explained *σ*^2^_*explained*_ = *σ*^2^_*tot*_ – *σ*^2^_*res*_. In the case where *σ*^2^_*res*_ > *σ*^2^_*tot*_, we have variance added by model 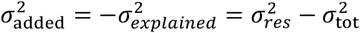.

##### Nucleotide importance score

We calculated nucleotide importance score (NIS) using ΔΔ*G*_37_ for mismatches, and Δ*G*_37_ for triloop, tetraloop, and bulges. For a dataset of *n* sequence variants with *n* corresponding energy values *y*_1_, …, *y*_*n*_, either Δ*G*_37_ or ΔΔ*G*_37_, we group the *n* data points into four groups based on the identity of the nucleotide of interest. For each of the four groups, we calculated the mean, 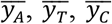, and 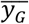. The grand mean 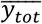 is the mean of *y*_1_, …, *y*_*n*_. The NIS is simply the between group sum of squares (SSB):

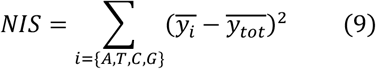

##### Pairwise interaction score

For a pair of nucleotides of interest, *M* and *N*, we compare two linear models that predict the energy value *Y*, model A with independence assumption and model B without. Model A is in the form of *M* + *N∼Y*, and model B *MN∼Y*. The pairwise interaction score (PIS) is the difference of *R*^2^ between the two linear models

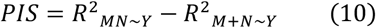

### Calculations with NUPACK

#### Melting temperature

Melting temperatures of DNA duplexes were calculated from the fraction of molecules unpaired. We first calculated structure free energies of the paired state at temperature T, Δ*G*_*paired,T*_, then got the equilibrium constant

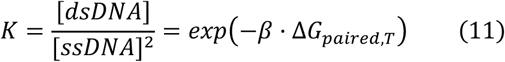

where *β* = 1/*RT* is the thermodynamic beta, [*dsDNA*] is the concentration of double-strand DNA, and [ssDNA] that of the single-strand DNA. At melting temperature, [*dsDNA*] = [*ssDNA*] = [*DNA*]/2 for distinguishable strands and [*dsDNA*] = [*DNA*]/4 for indistinguishable strands, where [*DNA*] is the total concentration of DNA strand species.

This gives 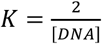 or 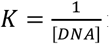 for distinguishable and indistinguishable strands respectively. Solving this equation with bisection gives us the melting temperature.

For DNA hairpins, melting temperature is independent of DNA strand concentration. The Boltzmann factor becomes:

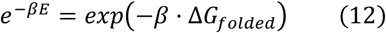

*T*_*m*_ is then simply the temperature when *dG*_*folded*_ = 0. Therefore, we get structure free energies Δ*G*_1_ and Δ*G*_2_ at a lower temperature *T*_1_ and a higher one *T*_2_, then calculate Δ*S* = −(Δ*G*_2_ − Δ*G*_1_)/(*T*_2_ – *T*_1_), Δ*H* = Δ*G*_1_ + *T*_1_ · Δ*S*, and *T*_*m*_ = Δ*H*/Δ*S*.

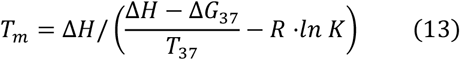

#### Data splitting for machine learning models

After filtering with the two-state criteria as described in “ΔG line fitting”, we splitted the clean Array Melt into train, validation, and test sets, stratified by variant type. We first splitted 5% of the data as the test set (1438 variants), then splitted 5% of the remaining data as the validation set (1315 variants), and 95% of the rest as the training set (24979 variants). We splitted the external 384-duplex dataset and Oliveira et al. dataset to a 1:1 split of validation and test data. The cloud lab UV melting data were used for correction between Array Melt and bulk solution and were not used for model testing to prevent circularity.

#### Linear models

##### Weighted linear regression model

We used a singular value decomposition (SVD) solver to solve the linear regression solutions, taking measurement error of each sequence into account. For a training dataset of *n* sequence variants with *n* corresponding Δ*H* or Δ*G*_37_ values *y*_1_, …, *y*_*n*_, and measurement uncertainty values 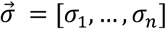, we have a *n* by *n*_*feature*_ feature count matrix *X* derived from the sequences. We then did SVD on matrix 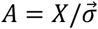 and get *U, S, V*. Any diagonal value in *S* that is smaller than a relative numeric threshold 10^−15^ was set to zero. Then the estimated parameter values for the features are 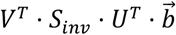, and the estimated errors for the features are 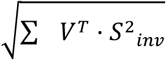, where 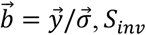 is *S* but taking 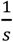 for all diagonal elements, and *S*^2^_*inv*_ is *S* but taking 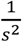 for all elements.

##### The NUPACK-compatible nearest-neighbor model

We trained the improved nearest-neighbor model on the Array Melt training data set, containing 24,971 of the 27,732 sequences that passed quality control and two-state criteria, and tuned the model on a validation set of 1,315 sequences. We first adjusted salt concentration to 1M Na^+^ to be compatible with NUPACK and other softwares, and then corrected systematic errors by linearly shifting Tm and ΔG, such that the medians of each construct type match those of NUPACK predictions. The underlying assumption is that NUPACK predictions have minimal bias but could be improved by explaining more variance. Improved parameters for NUPACK were fitted with weighted linear regression. We allowed an extra intercept parameter to adjust mean absolute error. From sequence-structure pairs in the training set, we extracted features of stacks and loops. Stacks include nearest-neighbor Watson-Crick pairs and terminal stack penalty, as in NUPACK; loops include the following:

1. Hairpin loops, defined as a subsequence *ϕ*_*i*:*j*_ where *i* · *j* is the closing base pair. We extracted “hairpin_mismatch” features as in NUPACK, 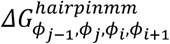, and introduced an intermediate “hairpin_loop_mid” feature, 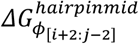 to account for sequence variations in the middle of hairpin loops for triloops and tetraloops. The final energy of hairpin is 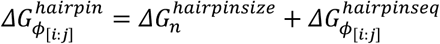 as in^20^, where

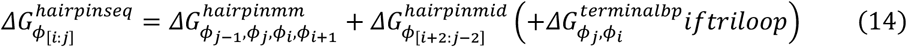
2. Interior loops, defined from 5’ to 3’ as two subsequences *ϕ*_[*i*:*d*]_ and *ϕ*_[*e*:*j*]_. Any other interior loop types were defined as in^20^, apart from the 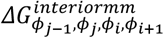, which were fitted separately instead of shared with terminal mismatches as in NUPACK, and that we introduced an intermediate parameter for the interaction between the two base pairs surrounding 1x1 mismatches 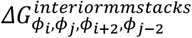.

This results in 178 features for *ΔH* and *ΔG* each. Parameters for hairpin and interior loop sizes were taken from NUPACK and clamped during fitting. Lookup tables of 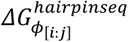 and 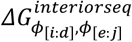 for “interior_1_1”, “interior_1_2”, and “interior_2_2” were calculated from the fitted parameters. All updated parameters were then written to a NUPACK-compatible json file.

Afterwards, we obtained parameters for 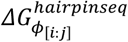 for tetraloops and triloops directly from the ‘2nd flanking’ construct types of either tetraloop or triloop parts of the library (**Fig. 1c**). These variants covered all possible sequences for hairpin loops of length 3 or 4, as well as the nearest-neighbor and next-nearest-neighbor closing pair sequences. We first regressed out the next-nearest-neighbor closing pair of the loops from measured ΔH and ΔG values, then aggregated the mean values for variants with identical hairpin loop and nearest-neighbor closing pair sequences. These aggregated values have the correct ΔΔH and ΔΔG relationships between each other but need a constant to be added to get the ΔH and ΔG values needed for the nearest-neighbor model. To solve this problem, we fitted two correction values *Δ*Δ*H* and Δ*ΔG* in kcal/mol. We searched for values that minimize the equation

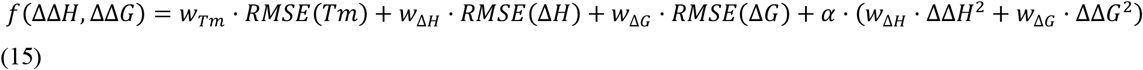

where *w*_*Tm*_ = 4, *w*_Δ*H*_ = 0.3, *w*_Δ*H*_ = 3.4 and *α* = 0.0001 are weights determined by validation on 100 sampled variants in the training set. Calculating this equation is computationally expensive, as it involves directly calling NUPACK in each iteration. We minimized it on 25 randomly sampled variants from the training set, repeating the process 4 times on different subsets using the BFGS algorithm implemented in the ‘scipy.optimize.minimize’ function, with parameters tol=1e-1 and maxiter=20. The resulting *Δ*Δ*H* and Δ*ΔG* values were mostly consistent between the runs (**Fig. S4g**). We took the median values from the 4 runs (ΔΔ*H* = −28 *kcal*/*mol*, ΔΔ*G* = −4 *kcal*/*mol*) and added them to the aggregated ΔΔ*H* and *Δ*Δ*G* values of hairpin loops.

Empirically, we found that model performance on the validation set was improved when we augmented the compact 178-parameter nearest-neighbor model with these curated hairpin loop parameters. This improvement might be due to the dominantly overrepresentation of ‘GAAA’ hairpin loops, which led to highly skewed data distribution and caused directly regressed hairpin loop parameters to be less accurate.

##### The richer parameter nearest-neighbor model

We trained the richer parameter model on the same Array Melt training data set as for the NUPACK-compatible nearest-neighbor model. For feature extraction, we extracted nearest-neighbor Watson-Crick pairs and terminal stack penalty as in NUPACK, and the following parameters for loops:

1. Hairpin loops. For triloops and tetraloops, 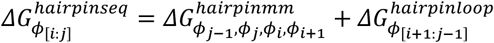, where 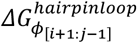 is the full sequence of the hairpin loop, in contrast to only the middle of loops in the NUPACK-compatible model. For hairpin loops longer than tetraloop, we applied a single loop sequence parameter 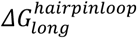. The final energy of hairpin is also 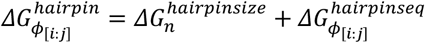.
2. Interior loops. For single and double bulges or single mismatches, the sequence-dependent parameter is fitted for each sequence, 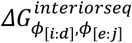. For double and triple mismatches, 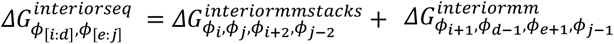.

### Deep learning models

#### Closest sequence lookup baseline model

We defined the distance between any two sequence-structure pairs by summing up the edit distance between the sequences and the dot-bracket secondary structure representation. For every query sequence-structure pair, we find the k closest sequence-structure pairs in the training set, whose distance-weighted sum becomes the predicted value for the query using python package *scipy*^39^. We found the best value for k using the held-out Array Melt validation data.

#### Graph neural network model

We implemented the graph neural network using python packages PyTorch and PyTorch Geometric^40^. The final model has 4 graph transformer layers with a 125-vec node attribute at each layer, stacked with dropout of 0.01 at each layer, a Set2Set module with 10 processing steps, and a 1 fully connected hidden layer with 128 channels and a dropout ratio of 0.49. We normalized ΔH and Tm of the training data to fall within 0 and 1 and used the sum of the root mean square error (RMSE) of both ΔH and Tm as the loss function and trained with Adam optimizer for 200 epoches. The model has 287,136 parameters. The validation model registry was monitored and stored on weight and biases (wandb.ai).

#### Using the NUPACK-compatible model

The parameter file for the NUPACK-compatible nearest-neighbor model is available in the supplementary material as ‘NUPACK_compatible_NN_model.json’. To use this file, download and point to it when using the python package NUPACK 4^41^:

my_model = nupack.Model(material=‘/path/to/parameter/file/NUPACK_compatible_NN_model.json’, …)

## Code availability

Code for array data preprocessing from raw sequencing and imaging data plus curve fitting could be found at https://github.com/GreenleafLab/array_analysis.

All subsequent analysis could be found at https://github.com/GreenleafLab/nnn_paper.

## Supplementary Figures

**Figure S1.**
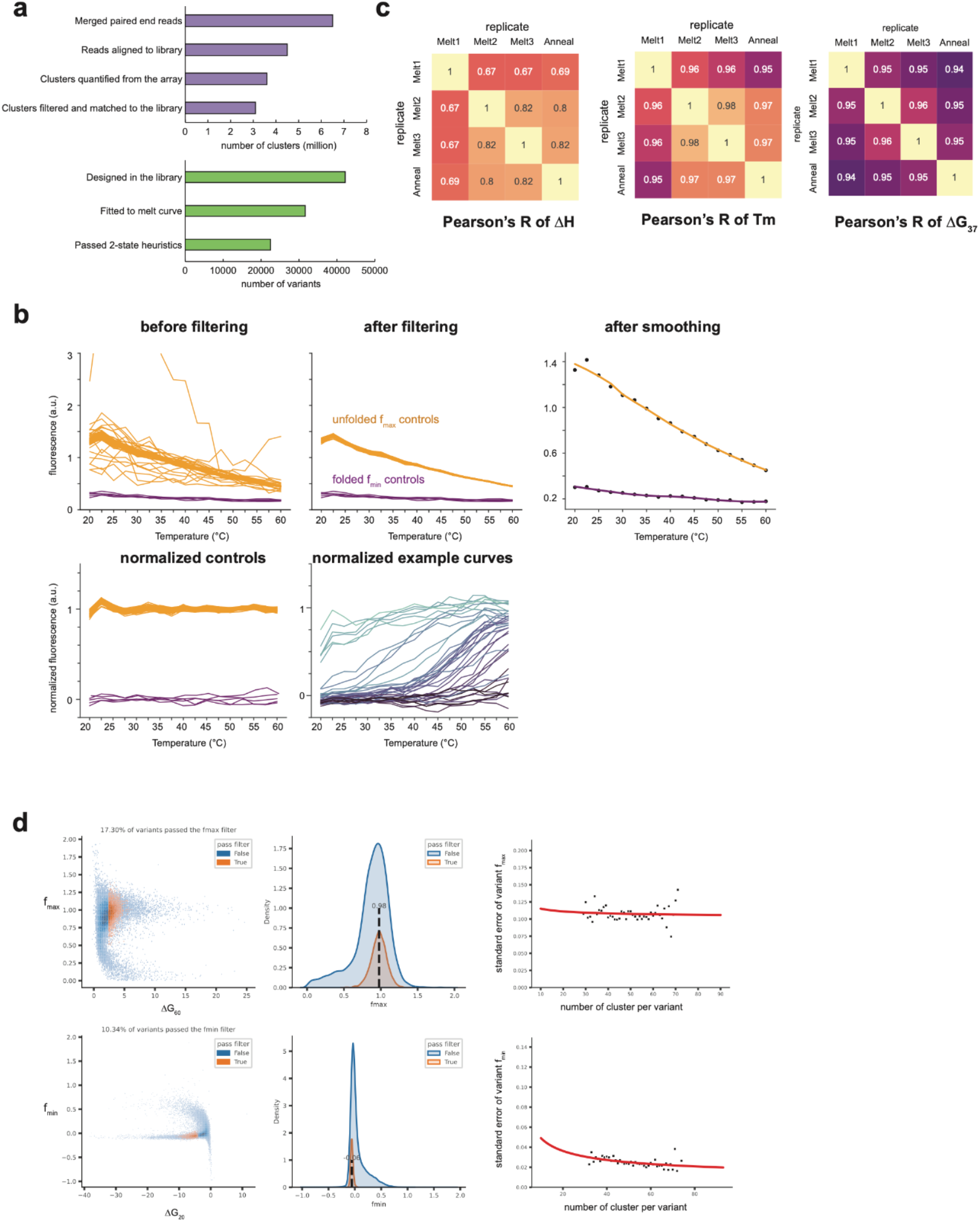
a. Number of measurements in a single Array Melt experiment. b. Pearson’s R of ΔH, Tm, and ΔG_37_ between replicates. c. Controls used to normalize fluorescence across different temperatures, including always unfolded constructs for f_max_ (maximum fluorescence) and folded constructs for f_min_ (minimum fluorescence). Some Watson-Crick curves are randomly selected to visualize after the normalization, colored by their expected ΔG_37_ values predicted by the nearest-neighbor model. d. Estimation of f_max_ and f_min_ after the initial round of curve fitting. Of all the fitted variants (blue), only a fraction that passed quality control filters (orange) were used to estimate f_max_ and f_min_ parameters used for the next round of refinement fitting. Uncertainties of f_max_ and f_min_ were corrected as a function of number of clusters per variant by curve fitting (right).

**Figure S2.**
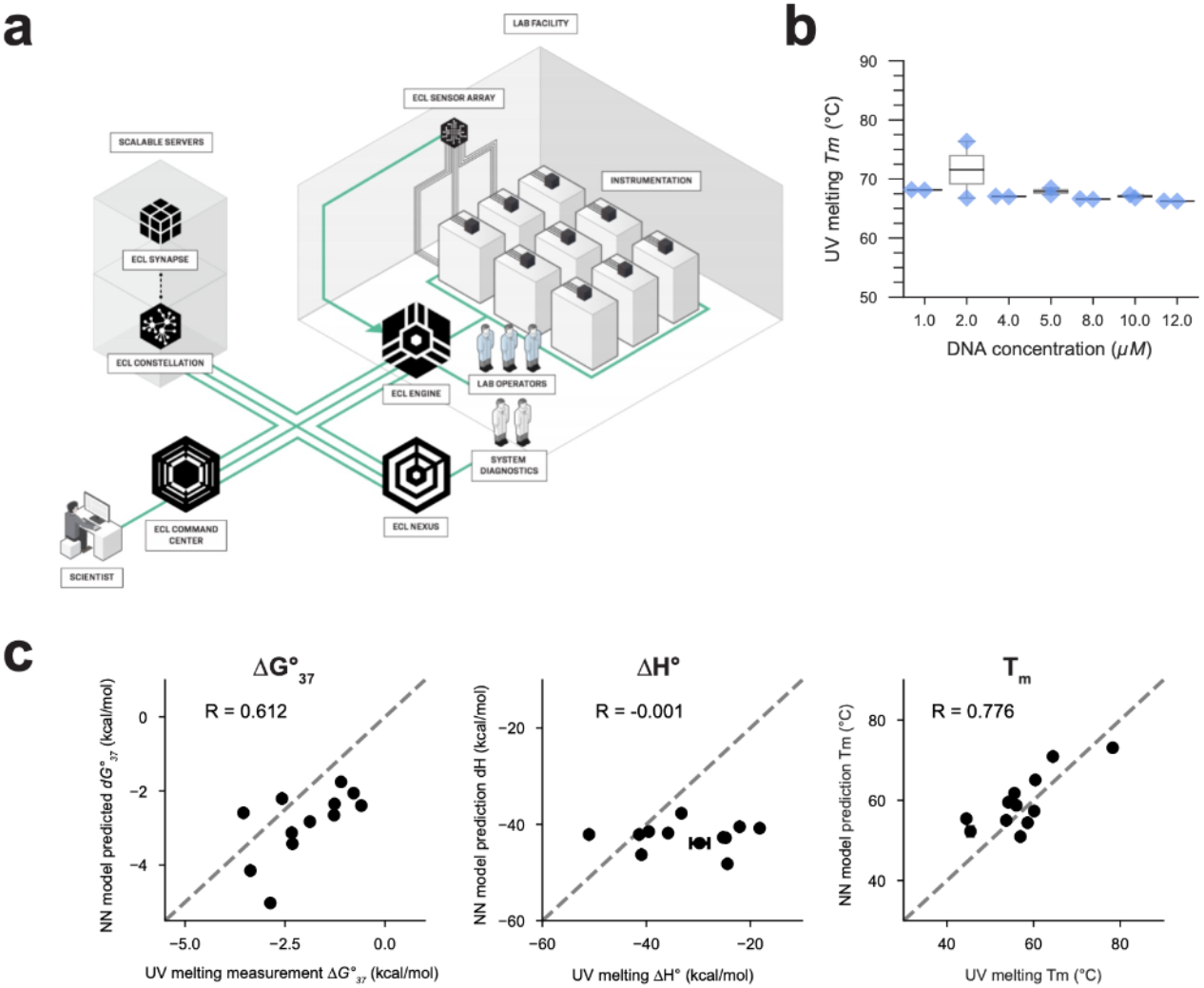
a. Schematic representation of Emerald Cloud Lab (ECL) workflow. Image from ECL with permission. b. UV melting measured Tm at different DNA hairpin oligo concentrations. Oligo used is WC68. c. Direct comparison of measured ΔG, ΔH and Tm between nearest-neighbor model prediction and UV melting data. Each datapoint is one hairpin variant.

**Figure S3.**
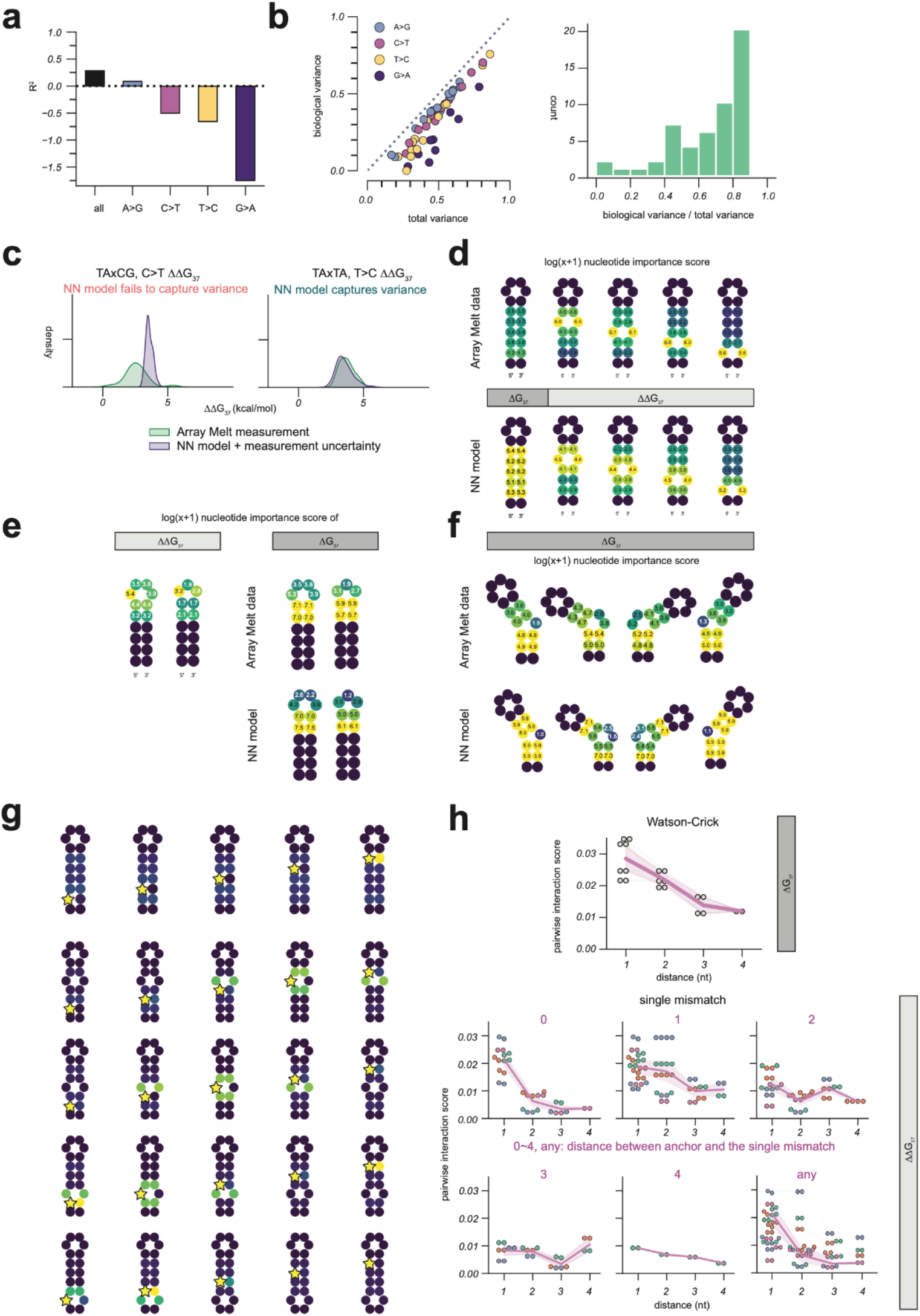
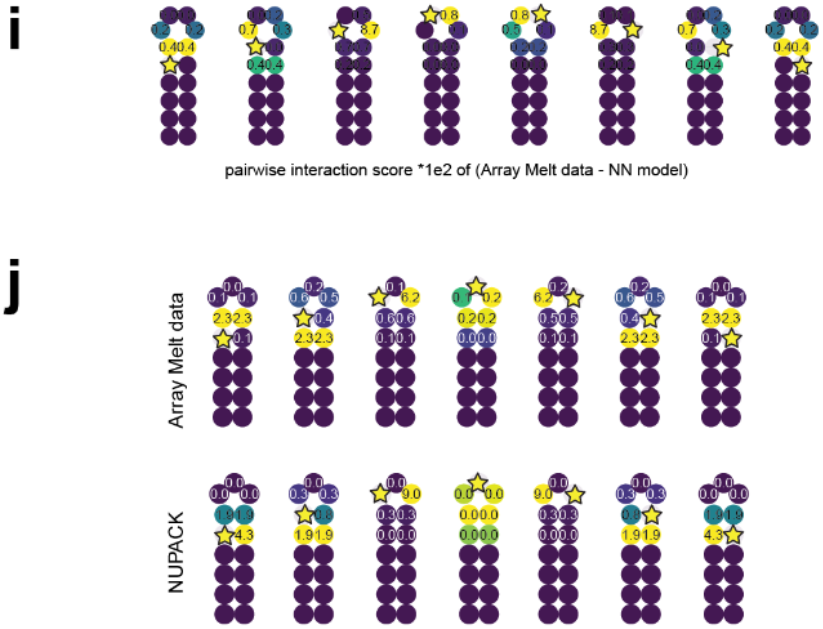
a. R^2^ values between Array Melt data and nearest-neighbor model predictions for all single mismatches, and for each mismatch type. b. Relationship of biological variance to total variance in Array Melt data. Each dot is a group of variants with identical mismatch type and adjacent nearest-neighbor base pairs as in **Fig. 3a**, colored by mismatched type. The biological/total variance ratio is shown on the right. c. Examples of when the nearest-neighbor model succeeds or fails to capture total variance in data. Each subplot is calculated from a group of variants with identical mismatch type and adjacent base pairs. Green is the histogram of Array Melt measurements, and purple is what is expected from the nearest-neighbor model prediction with measurement uncertainty. d. Comparison of nucleotide importance scores calculated from Array Melt data and nearest-neighbor model predictions for Watson-Crick and single mismatch variants. e. Comparison of nucleotide importance scores calculated from either ΔΔG37 or ΔG_37_, and on Array Melt data or nearest-neighbor model predictions, for tetraloops and triloops. f. Comparison of nucleotide importance scores calculated from Array Melt data and nearest-neighbor model predictions of bulge variants. g. Pairwise interaction scores calculated from nearest-neighbor model predictions for Watson-Crick and single mismatch variants. h. Pairwise interaction scores as a function of distance between the two nucleotides in the pair, for Watson-Crick variants or single mismatch variants. Those for the latter are splitted by the distance from the anchor point to the single mismatch. i. Difference between pairwise interaction scores from Array Melt data and nearest-neighbor model prediction. j. Comparison of nucleotide importance scores calculated from Array Melt data and nearest-neighbor model predictions of triloop variants.

**Figure S4.**
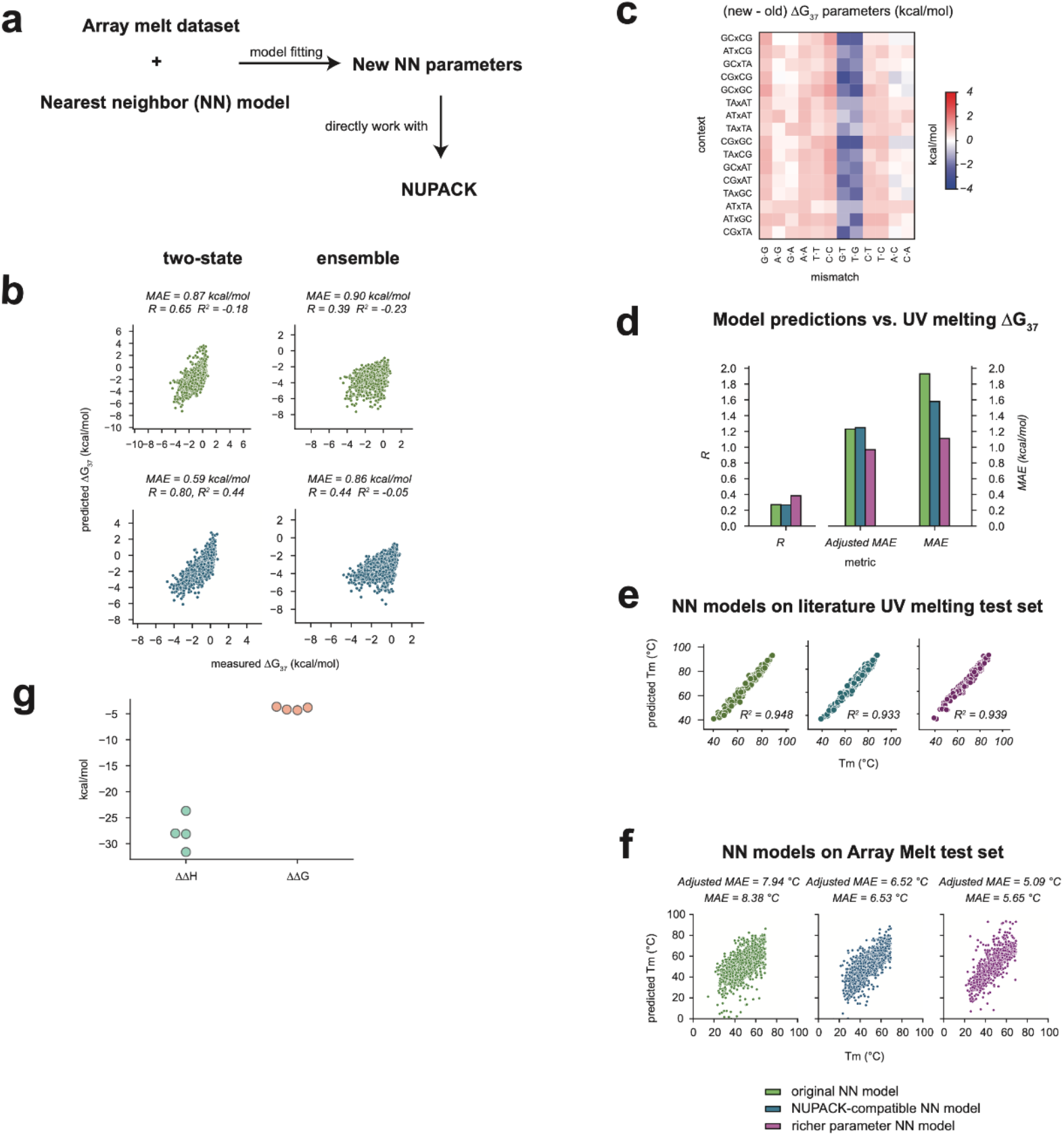
a. Schematics of the NUPACK-compatible nearest-neighbor model. b. Difference between the new and original *dna04* single mismatch parameters. c. Comparison of two-state and ensemble model predictions from NUPACK against Array Melt measurements. d. Comparison of *dna04*, NUPACK-compatible, and richer parameterization nearest-neighbor model predictions with cloud lab UV melting measurements of randomly designed DNA hairpins. e. Performance of the original, NUPACK-compatible, and richer parameter nearest-neighbor models on literature UV melting data used to derive the original nearest-neighbor model. f. Performance of the original, NUPACK-compatible, and richer parameter nearest-neighbor model on Array Melt test set for Tm. g. Fitted ΔΔH and ΔΔG values for ‘hairpin_tetraloop’ and ‘hairpin_triloop’. Each dot is from one individual optimization run.

**Figure S5.**
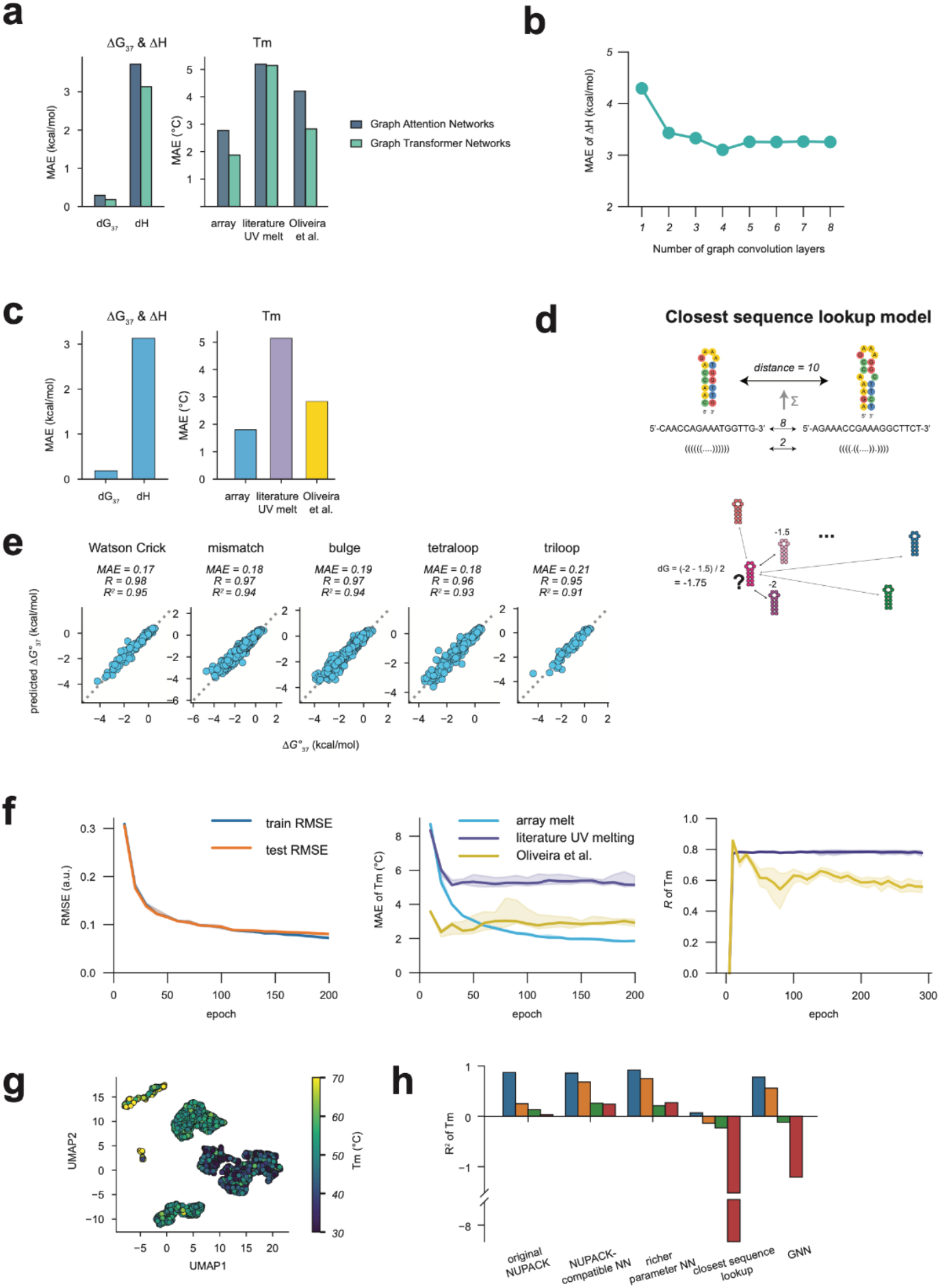
a. Mean absolute error of GNNs with either Graph Attention Network or Graph Transformer Network architecture at the graph convolutional layer. b. Mean absolute error of enthalpy on Array Melt data as a function of the number of graph convolution layers in the GNN. c. Mean absolute error of melting temperature on Array Melt data as a function of percentage of training data used to fit the GNN. d. Schematic of the baseline closest sequence lookup model. e. GNN predicted ΔG_37_ vs. held out Array Melt, splitted by type of variant. f.The objective function of GNN (RMSE of normalized ΔH and Tm), mean absolute error of Tm, and Pearson’s R of Tm across the GNN training process. g. UMAP of learned embeddings of test data, colored by their predicted Tm. h. Benchmark of all models on the orthogonal Oliveira et al. dataset as in **Fig. 5** but plotted by model. Bars for each model are for no mismatch, one mismatch, two mismatches and three mismatches, respectively.

## Supplementary Notes

### Adjusted mean absolute error

Nearest-neighbor models inherently have parameterization degeneracy, meaning that adding a constant to one group of parameters and subtracting from another could lead to the same prediction. This degeneracy can result in a prediction bias, but its toll on prediction accuracy could be mitigated by applying a constant offset to align the predictions with the true values. This constant offset preserves the relative ΔΔ*G* values but not the absolute ΔG values. For instance, in the Array Melt test set, NUPACK with the original parameter set has a ΔG_37_ bias of -0.71 kcal/mol, and our NUPACK-compatible model parameter set has a bias of -0.42 kcal/mol. After subtracting this bias to align the predictions with the true values, the MAE of ΔG_37_ was reduced from 1.07 to 0.87 kcal/mol for the original NUPACK model, and from 0.67 to 0.59 kcal/mol for our NUPACK-compatible model (**Table 1**). This adjustment demonstrated that, with the presence of this bias, the accuracy of predicted ΔΔ*G* values between different sequence-structure pairs is generally higher than that of the predicted Δ*G* values compared to the measured ground truth. Despite this model bias, we argue that the accuracy of nearest-neighbor models is acceptable in practice, particularly for applications that primarily rely on relative ΔΔ*G* values rather than absolute Δ*G* values. This is because many applications, such as using dynamic programming to find the minimum free energy structure, depend on the relative ΔΔ*G* values between different candidate secondary structures.

Given this context, we find it reasonable to report both the raw and the adjusted MAE model metrics. The adjusted metrics are calculated by first centering the prediction values to match the mean of the true values, thereby providing a more informative estimate of the relative errors for a given model.

